# Uncovering temporal structure in hippocampal output patterns

**DOI:** 10.1101/242594

**Authors:** Kourosh Maboudi, Etienne Ackermann, Brad Pfeiffer, David Foster, Kamran Diba, Caleb Kemere

**Author notes:** **For correspondence:** (KD); (CK). These authors contributed equally to this work (Kourosh Maboudi, Etienne Ackermann, Kamran Diba, Caleb Kemere).

## Abstract

The place cell activity of hippocampal pyramidal cells has been described as the cognitive map substrate of spatial memory. Replay is observed during hippocampal sharp-wave ripple-associated population burst events and is critical for consolidation and recall-guided behaviors. To present, population burst event (PBE) activity has been analyzed as a phenomenon subordinate to the place code. Here, we use hidden Markov models to study PBEs observed during exploration of both linear mazes and open fields. We demonstrate that estimated models are consistent with temporal replay sequences and that the latent states correspond to a spatial map of the environment. Moreover, we demonstrate the identification of hippocampal replay without recourse to the place code, using only PBE model congruence. These results suggest that downstream regions may rely on PBEs to form a substrate for memory. Additionally, by forming models independent of animal behavior, we lay the groundwork for studies of non-spatial memory.

## Introduction

Large populations of neurons fire in tandem during hippocampal sharp-waves and their accompanying CA1 layer ripple oscillations (*Buzsáki, 1986*). By now, multiple studies have shown that during many sharp-wave ripple-associated population burst events (PBEs), hippocampal “place cells” (*O’Keefe, 1976*) fire in compressed sequences that reflect the firing order determined by the sequential locations of their individual place fields (*Diba and Buzsáki, 2007; Foster and Wilson, 2006; Lee and Wilson, 2002; Nádasdy et al., 1999*). While the firing patterns during active exploration are considered to represent the brain’s global positioning system and provide a substrate for spatial and episodic memory, instead it is the synchronized activity during PBEs that is most likely to affect cortical activity beyond the hippocampus (*Buzsáki, 1989; Carr et al., 2011; Diekelmann and Born, 2010; Siapas and Wilson, 1998*). Likewise, widespread activity modulation is seen throughout the brain following these sharp-wave ripple population bursts (*Logothetis et al., 2012*).

The literature on PBEs has largely focused on developing templates of firing patterns during active behavior and evaluating the extent to which these templates’ patterns are reprised during subsequent PBEs. But what if the fundamental mode of the hippocampus is not the re-expression of place fields, but rather the PBE sequences during sharp-wave ripples (SWRs)? PBE sequences are enhanced during exploration of novel environments (*Cheng and Frank, 2008; Foster and Wilson, 2006*), they presage learning-related changes in place fields (*Dupret et al., 2010*), and appear to be critical to task learning (*Ego-Stengel and Wilson, 2010; Girardeau et al., 2009; Jadhav et al., 2012*). Here, we examine the information provided by CA1 and CA3 pyramidal neurons, the output nodes of the hippocampus, through the looking glass of PBE firing patterns.

We developed a technique to build models of PBE sequences strictly outside of active exploration and independent of place fields and demonstrate that this nevertheless allows us to uncover spatial maps. Furthermore, these models can be used to detect congruent events that are consistent with replay but without any explicit place cell template. Our technique therefore provides new possibilities for evaluating hippocampal output patterns in single-trial and other fast learning paradigms, where a reliable sequential template pattern is not readily available. Overall, our work suggests that a sequence-first approach can provide an alternative view of hippocampal activity that may shed new light on how memories are formed, stored, and recalled.

## Results

### Awake population burst events

We first used previously-recorded activity of large numbers of individual neurons in areas CA1 and CA3 of the dorsal hippocampus as rats navigated linear mazes (*Diba and Buzsáki, 2007*) for water reward (linear track: *n* = 3 rats, *m* = 18 sessions). Using pooled multiunit activity, we detected PBEs during which many neurons were simultaneously active. The majority of these events (55.0% ± 20.1% sd), occurred when animals paused running to obtain reward, groom, or survey their surroundings (*Buzsáki et al., 1983*), and were accompanied by SWR complexes, distinguished by a burst of oscillatory activity in the 150–250 Hz band of the CA1 local field potential (LFP). Because we are interested in understanding internally generated activity during PBEs, we excluded periods of active behavior (running speed > 5 cm/s), ensuring that theta sequences would not bias our results. We did not add any other restrictions on behavior, LFPs, or the participation of place cells. We found that these non-theta PBEs occupied an average of 1.8% of the periods during which animals were on the linear track (16.9 ± 15.1 s of 832.6 ± 390.5 s), in comparison to the 34.8% of time animals were running (running speed > 10 cm/s) on the track (254.4 ± 106.6 s of 832.6 ± 390.5 s).

### Learning hidden Markov models from PBE data

Activity during PBEs is widely understood to be internally-generated in the hippocampal-entorhinal formation, and likely to affect neuronal firing in downstream regions (*Buzsáki, 1989; Chrobak and Buzsáki, 1996; Logothetis et al., 2012; Yamamoto and Tonegawa, 2017*). Given the prevalence of PBEs during an animal’s early experience, we hypothesized that the neural activity during these events would be sufficient to train a machine learning model of sequential patterns—a hidden Markov model—and that this model would capture the relevant spatial information encoded in the hippocampus independent of exploration itself. In our hidden Markov models (HMMs), the unobserved latent variable represents the temporal evolution of a memory trace which is coherent across the CA1 ensemble. We used a discretized latent space with a fixed number of states to simplify model fitting. The parameters of our model include the initial state distribution (the probability for sequences to start in each state), the state transition model (the probability that the CA1 ensemble will transition from a start state to a destination state in the next time bin), and the observation model (predicted activity of each excitatory neuron within the CA1 ensemble for a given state). In prior work using HMMs to model neural activity, a variety of statistical distributions have been used for observation models (*Chen and Wilson, 2017; Chen et al., 2012,2014; Deppisch et al., 1994; Kemere et al., 2008; Radons et al., 1994*). We opted for the Poisson distribution to minimize the number of parameters per state and per neuron. We used the standard iterative expectation-maximization (EM) algorithm (*Rabiner, 1989*) to learn the parameters of an HMM which represented binned PBE data (20 ms bins). *Figure 1* depicts the resultant state transition matrix and observation model for an example linear-track session.

**Figure 1.**
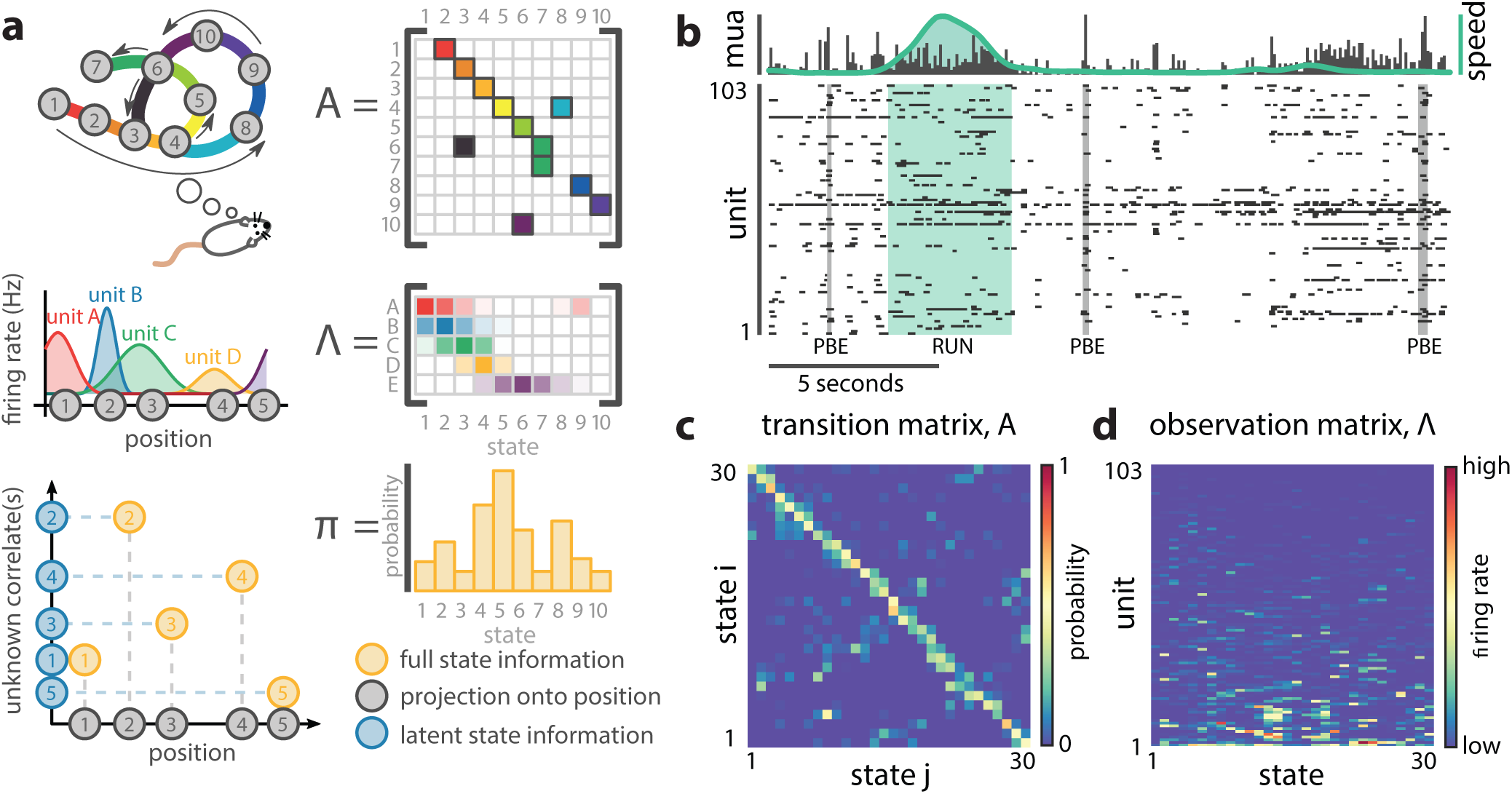
A hidden Markov model of ensemble activity during population burst events. **a.** A cartoon illustration of our HMM and how it relates to imagined position-specific states (top row, state transition matrix), unit firing activity (middle row, observation matrix), and additional unobserved state information (bottom row, state information representation and prior probabilities overstates). **b.** Examples of three PBEs and a run epoch. **c.** The transition matrix models the dynamics of the unobserved-but-coherent internally-generated state. The sparsity and banded-diagonal shape are suggestive of sequential dynamics. **d.** The observation model of our HMM is a set of Poisson probability distributions (one for each neuron) for each hidden state. Looking across columns (states), the mean firing rate is typically elevated for only a few of the neurons and individual neurons have elevated firing rates for only a few states. Figure 1-Figure supplement 1. Hidden Markov models capture state dynamics beyond pairwise co-firing.

Parameter learning using the EM algorithm on limited data can yield apparent structure even if there is none. Using separate training and test-data sets (cross-validation) mitigates over-fitting to training data, but it is still possible for the cross-validated goodness-of-fit to increase with training without any underlying dynamics, e.g., if groups of neurons tend to activate in a correlated fashion. We wanted to understand whether the models we learned reflected underlying state dynamics of memory traces beyond pairwise co-firing. To answer this question, we cross-validated the model against both real “test” data and against surrogate “test” data derived from shuffling each PBE in two ways: one in which the binned spiking activity was circularly permuted across time for each neuron independently of the other neurons (“temporal shuffle”, which removes co-activation), and one in which the order of the binned data was scrambled coherently across all neurons (“time swap”, which maintains co-activation). Note that the second shuffle preserves any pairwise correlations while removing the order of any sequential patterns that might be present. Using five-fold cross-validation, we compared learned models against both actual and surrogate test data and found that the model likelihood was significantly greater for real data (vs. temporal shuffle, *p* < 0.001, vs. time swap, *p* < 0.001, *n* = 18 sessions, Wilcoxon signed-rank test, *Figure 1-Figure Supplement 1*).

### What do the learned model parameters tell us about PBEs?

To begin to understand what structure we learn from PBE activity, we compared our HMMs (trained on real data) against models trained on multiple different surrogate datasets (***Figure 2***a,b). These surrogate datasets were obtained from actual data following: 1) temporal shuffles and 2) time swaps, as above, and 3) by producing a surrogate PBE from independent Poisson simulations according to each unit’s mean firing rate within the original PBE. First, we quantified the sparsity of the observation model. We found that actual data yielded mean firing rates which were highly sparse (***Figure 2***d), indicating that individual neurons were likely to be active during only a small fraction of the states. Next, we investigated the transition matrices. Strikingly, the actual data yielded models in which the state transition matrix was more sparse than in each of the surrogate counterparts (*p* < 0.001, ***Figure 2***c), reflecting that intricate yet reliable details are captured by the HMMs. Using measures adopted from graph theory, we simulated paths through state space generated by these transition matrices, and found that this increased sparsity accompanied longer trajectories (*Figure 2-Figure Supplement 3*) through the state space of the model. Thus, the state transition matrices we learn are suggestive of dynamics in which each sparse state is preceded and followed by only a few other, in turn, sparse states, providing long sequential paths through state space-consistent with spatial relationships in the environment in which the animal was behaving, but generated from PBEs. The increased sparsity of the observation model and transition matrix in the example session was representative of a significant increase over all remaining sessions (*p* < 0.05, *n* = 18 sessions, Wilcoxon signed-rank tests, ***Figure 2***e,f).

**Figure 2.**
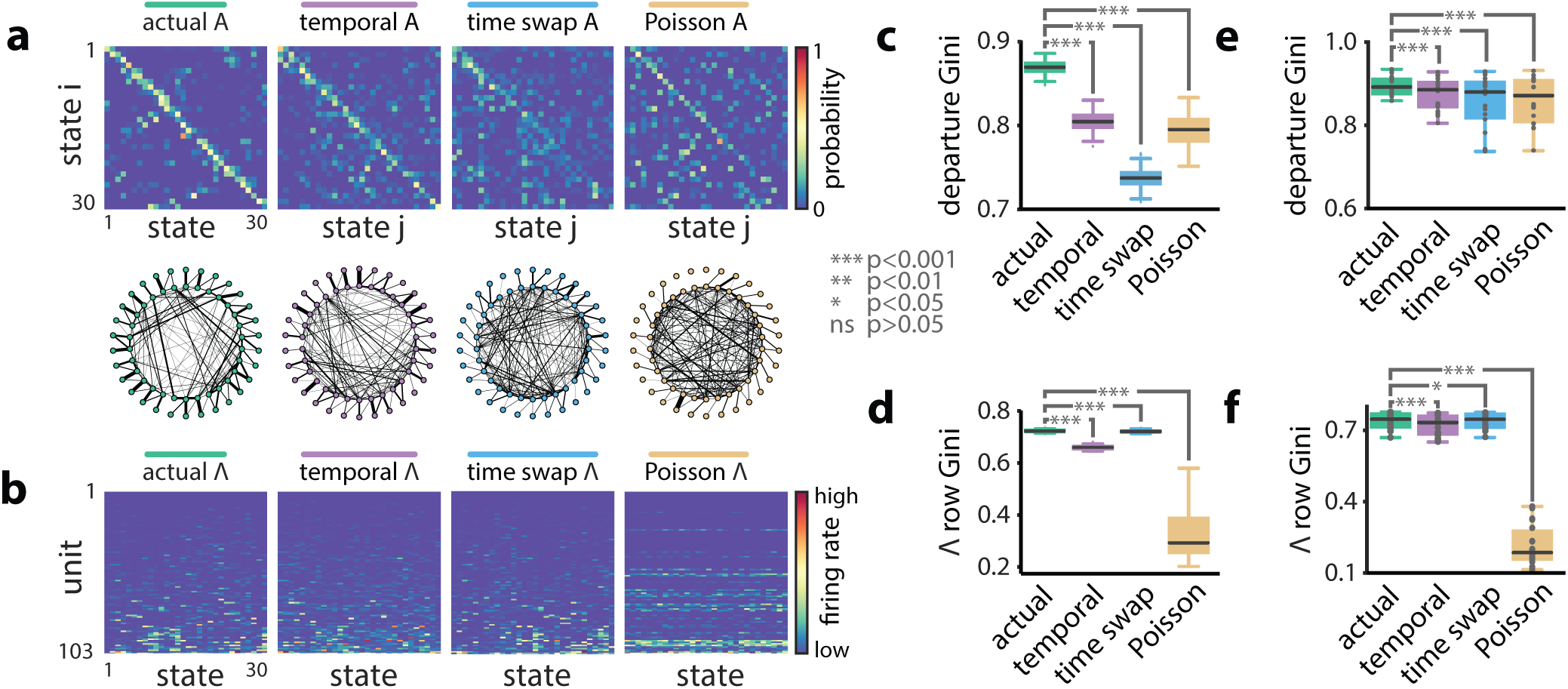
Models of PBE activity are sparse. We trained HMMs on neural activity during PBEs (in 20 ms bins), as well as on surrogate transformations of those PBEs. **a.** (top) The transition matrices for the actual and surrogate PBE models with states ordered to maximize the transition probability from state *i* to state *i* + 1. (bottom) Undirected connectivity graphs corresponding to the transition matrices. The nodes correspond to states (progressing clockwise, starting at the top) and nodes on the same radius correspond to the same state. The weights of the edges are proportional to the transition probabilities between the nodes (states). The transition probabilities from state i to every other state except *i* + 1 are shown in the interior of the graph, whereas transition probabilities from state *i* to itself, as well as to state *i* + 1 are shown between the inner and outer rings of nodes. **b.** The observation matrices for actual and surrogate PBE models show the mean firing rate for each neuron in each state. For visualization, neurons are ordered by their firing rates. **c.** We quantified the sparsity of transitions from one state to all other states using the Gini coefficient of rows of the transition matrix. For the example session in **a.**, actual data yielded sparser transition matrices than shuffles. **d.** The observation models learned from actual data are significantly more sparse than those learned after shuffling. This implies that as the hippocampus evolves through the learned latent space, each neuron is active during only a few states. **e.** Transition matrix sparsity and **f.** Observation model sparsity with corresponding shuffle data for all sessions/animals. (***: *p* < 0.001, *: *p* < 0.05; single session comparisons: *n* = 250 realizations, Welch’s t-test; aggregated comparisons - *n* = 18 sessions, Wilcoxon signed-rank test). Figure 2-Figure supplement 1. PBE model states typically only transition to a few other states. Figure 2-Figure supplement 2. Each neuron is active in only a few model states. Figure 2-Figure supplement 3. The sparse transitions integrate into long sequences through the state space.

These observations indicate that PBEs inform an HMM about extant spatial relationships within the environment. So, next we asked how the firing patterns of neurons during actual behavior project into the learned latent spaces. To observe the evolution of the latent states during behavior, we used our model and the forward-backward algorithm to decode neural activity in 100 ms bins during epochs that display strong theta oscillations (exclusive of PBEs) when rats were running (speed > 10 cm/s; see Methods). If the learned model was distinct from ensemble patterns during behavior, we might expect the resulting state space probability distributions at each point in time to be randomly spread among multiple states. Instead, we found distributions that resembled sequential trajectories through the latent space (***Figure 3***a) in parallel with the physical trajectories made by the animal along the track, further demonstrating that the latent state dynamics learned from PBEs corresponds to an internalized model of physical space.

**Figure 3.**
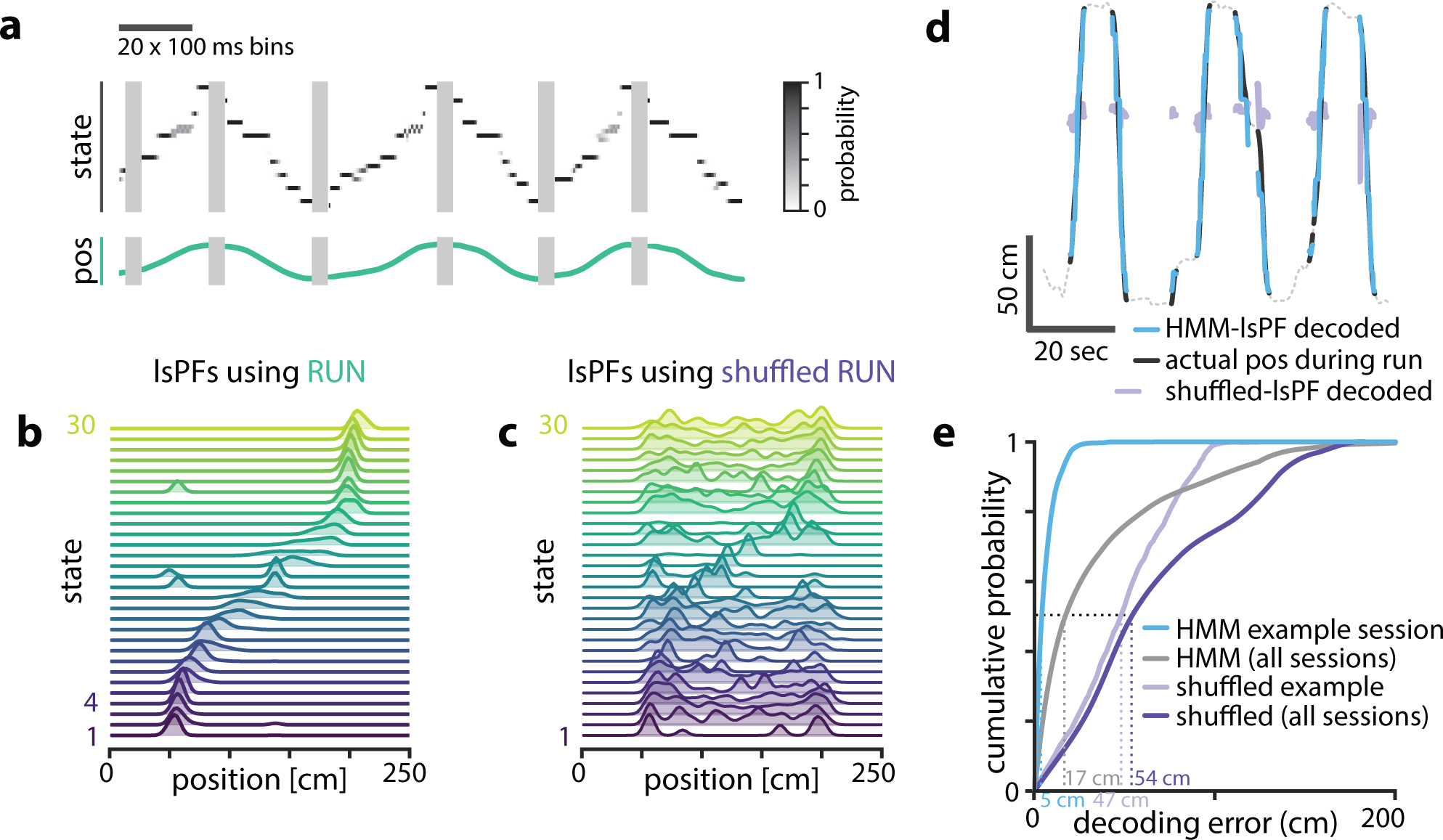
Latent states capture positional code. **a.** Using the model parameters estimated from PBEs, we decoded latent state probabilities from neural activity which was observed during periods in which the animal was running. An example shows the trajectory of the decoded latent state probabilities during 6 sequences of track running. **b.** Mapping latent state probabilities to associated animal positions yields latent-state place fields which describe the probability of each state for every position on the track. **c.** Shuffling the position associations yields uninformative state mappings. **d.** For an example session, position decoding through the latent space gives significantly better accuracy than decoding using the shuffled virtual tuning curves. **e.** The distribution of position decoding accuracy over all sessions (*n* = 18) was significantly greater than chance. (*p* < 0.001). Figure 3-Figure supplement 1. Latent states capture positional code over wide range of model parameters.

To better understand the relationship between the latent space and physical space, we used the latent state trajectories decoded during running with their corresponding positions to form an estimate of the likelihood as a function of location on the track (see Methods). These “latent-state place fields” (lsPFs, ***Figure 3***b) in many ways resembled neuronal place fields and similarly tiled the extent of the track. This spatial localization went away when we re-estimated the lsPFs with shuffled positions (***Figure 3***c). To quantify how informative the latent states were about position, we used the lsPF to map decoded state sequences to position during running periods (***Figure 3***d). In our example session, decoding through the latent space resulted in a median accuracy of 5 cm, significantly greater than the 47 cm obtained from shuffled lsPFs (*p* < 0.001, Wilcoxon signed-rank test, ***Figure 3***d). When we evaluated decoding error over our full set of sessions, we observed a similar result (*p* < 0.001, Wilcoxon signed-rank test, ***Figure 3***e, *Figure 3-Figure Supplement 1*). As our method required discretizing the state space, a potential caveat is that the number of latent states is a relevant parameter, which we arbitrarily chose to be 30. However, latent-state place fields were informative of position over a wide range of values of this parameter (*Figure 3-Figure Supplement 1*).

### HMM-congruent PBEs capture sequence replay

We and others have previously described how the pattern of place cell firing during many PBEs recapitulates the order in which they are active when animals run on the track (***Figure 4***a). We employed the versatile and widely-used Bayesian decoding method to ascribe a replay score to sequential patterns during PBEs. Briefly, for each PBE, we used place-field maps to estimate a spatial trajectory (an *a posteriori* distribution of positions) in 20 ms bins. We generated surrogate data via a column-cycle shuffle of the *a posteriori* distributions during PBEs. The real and surrogate trajectories were scored (see Methods), and we defined replay events as those for which the score of the actual trajectory was larger than a threshold fraction of the null distribution generated by the surrogate scores. Using this approach, we found that 57% of PBEs (1064 of 1883) were identified as replay beyond a threshold of 99% (median across datasets 54.2%, IQR = 32.8–61.0%, *Figure 4-Figure Supplement 1*). Thus, as has been reported many times (*Davidson et al., 2009; Diba and Buzsáki, 2007; Foster and Wilson, 2006; Karlsson and Frank, 2009*), only a fraction of PBEs (but many more than expected by chance) represent statistically significant replay. Given that we use all PBEs for model learning and our models capture the structure of the environment and the patterns expressed by place cells during exploration, we were interested in understanding whether we could also use our latent-space models to find these replay events. Indeed, for many events when we decode trajectories through state space, they resemble the sequential patterns observed when we decode position using Bayesian techniques and the place cell map (***Figure 4***b, left). However, given previous evidence for replay of environments not recently experienced (*Gupta et al., 2010; Karlsson and Frank, 2009*), we hypothesized that some PBEs might contain ensemble neural activity which is unstructured and thus unrelated to the learned model, and that these would correspond to the “non-replay” events found using traditional methods.

**Figure 4.**
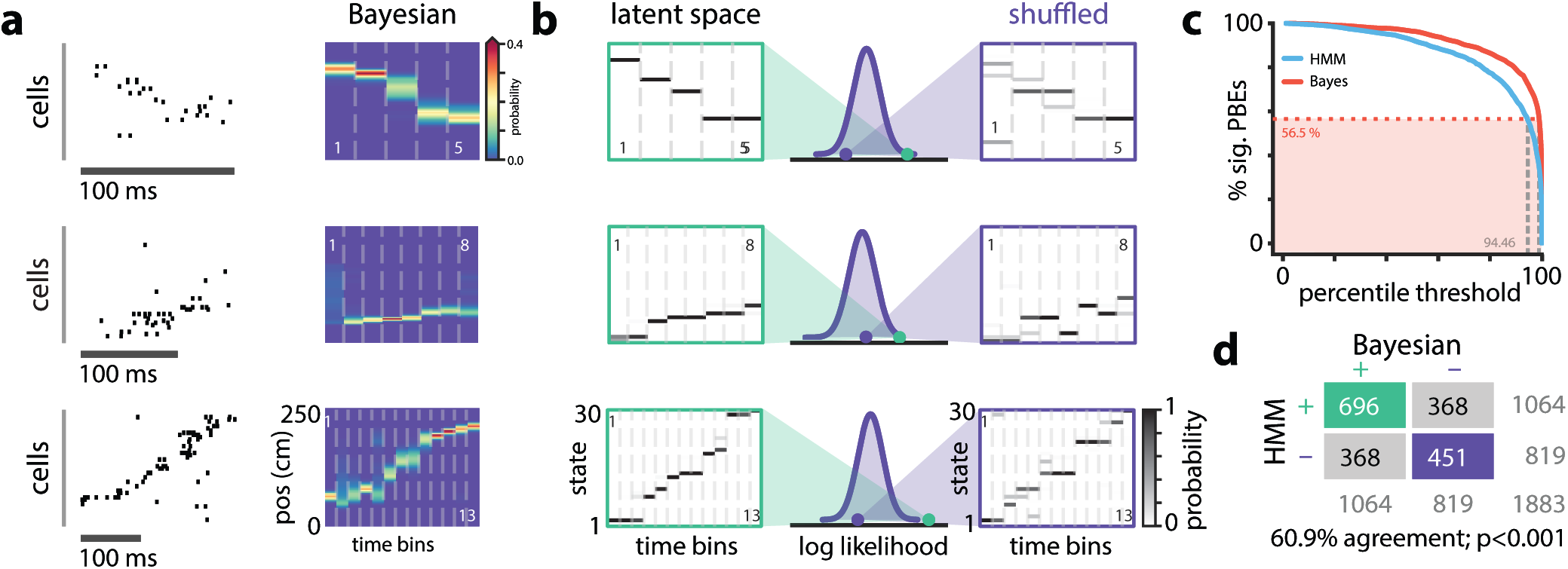
**a.** Example PBEs decoded to position using Bayesian decoding. **b.** (left) Same examples decoded to the latent space using the learned HMM. (right) Examples decoded after shuffling the transition matrix, and (middle) the sequence likelihood using actual and shuffled models. **c.** Effect of significance threshold on the fraction of events identified as replay using Bayesian decoding and model congruent events using the HMM-approach. **d.** Comparing Bayesian and model-congruence approaches for all PBEs recorded, we find statistically significant agreement in event identification (60.9% agreement, *n* = 1883 events from 18 sessions, *p* < 0.001, Fisher’s exact test two sided). Figure 4-Figure supplement 1. Number of significant PBEs.

To assess how well the pattern of ensemble activity during individual PBEs related to the overall state-space model learned from PBE activity (“congruence”), we developed a statistical approach for identifying the subset of strongly structured PBEs. Specifically, rather than comparing real and surrogate PBEs, we compared the goodness-of-fit for each event to a null distribution generated via a computationally-efficient manipulation of the transition matrix of the model (***Figure 4***b). We row-wise shuffle the non-diagonal elements of the transition matrix to assess whether an individual PBE is a more ordered sequence through state space than would be expected by chance. Maintaining the diagonal avoids identifying as different from chance sequences which consist of few states. As described above, the fraction of events identified as replay using Bayesian decoding is strongly tied to how the null-distribution is generated (i.e., what shuffle is used) and the value of the significance threshold arbitrarily chosen to be 90%, 95%, or 99% of shuffles in different reports. When we combined across datasets, we found that our transition matrix shuffle yielded a null distribution for which a 99% confidence interval identified slightly fewer PBEs as significant than the column-cycle shuffle did for Bayesian decoding (***Figure 4***c). To make a principled comparison of Bayesian- and HMM-based replay detection schemes, we fixed the Bayesian-based significance threshold at 99% but selected the significance threshold for the HMM-congruence null distribution so that the fraction of replay events detected would be the same between the two schemes. Following this approach, we found that model-congruent/incongruent PBEs largely overlapped with the replay/non-replay events detected using Bayesian decoding of the place cell map (***Figure 4***d). Thus, using only the neural activity during PBEs, without access to any place cell activity, we are remarkably able to detect the sequential patterns typically described as “replay” based only on their consistency with the structure of other PBE activity.

There were, however, also differences between the Bayesian and HMM-congruent approaches, including events that reached significance in one but not the other formalism. We wanted to understand where and why these approaches differed in identifying significant sequences. When we examined individual PBEs, we found sequences for which both Bayesian and model-congruence replay detection approaches appeared to malfunction (***Figure 5***a). This was not a failure of the choice of significance threshold, as for both techniques we found what appeared to be false-negatives (patterns which looked like replay but were not significant) as well as false-positives (patterns which looked noisy but were identified as replay). Thus, in order to quantitatively compare the two approaches, we asked eight humans to visually examine all the PBEs in our database. They were instructed to label as replay PBEs in which the animal’s Bayesian decoded position translated sequentially without big jumps (*Silva et al., 2015*, see Methods).

**Figure 5.**
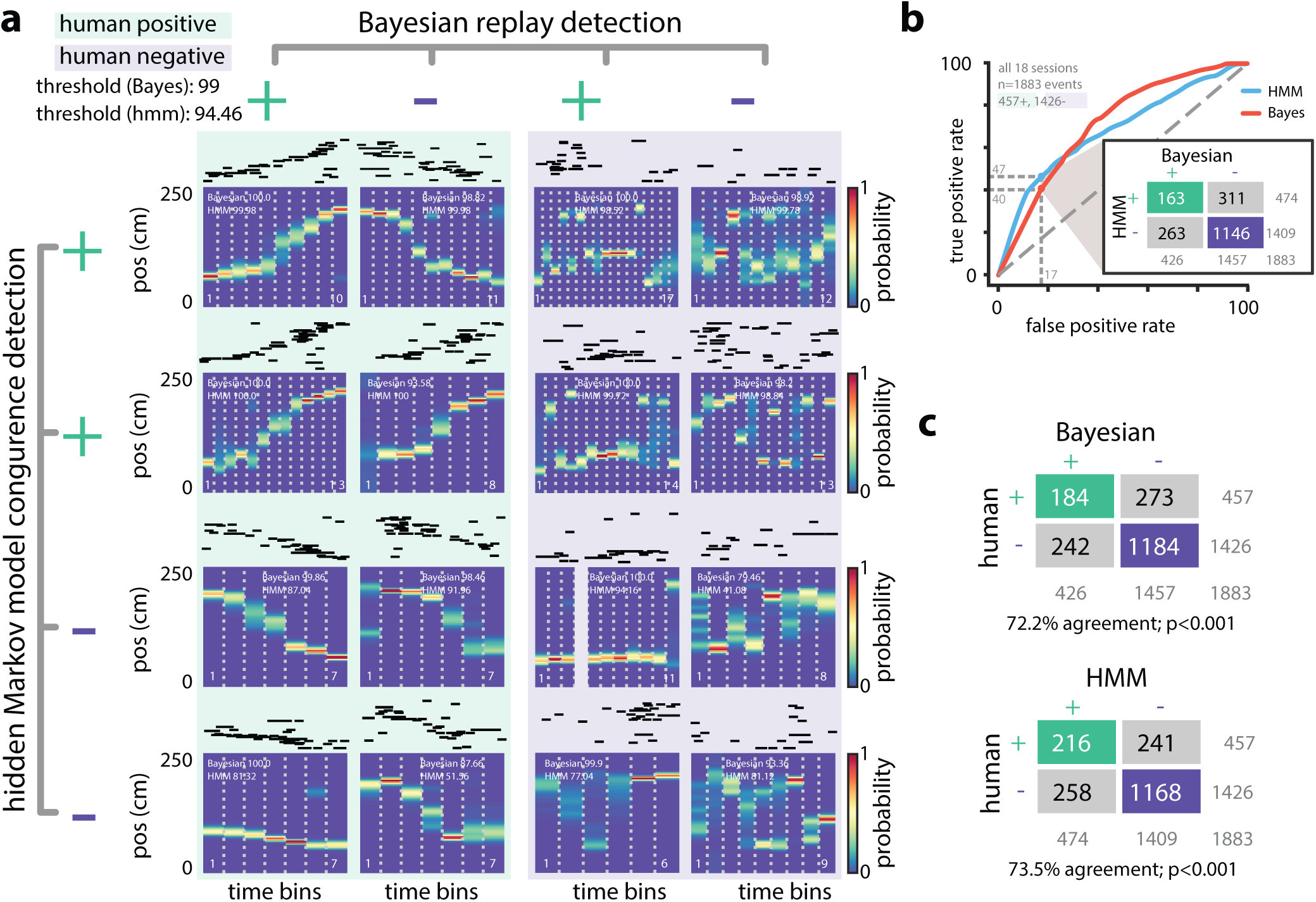
**a.** Eight examples from one session show that Bayesian decoding and HMM model-congruence can differ in labeling of significant replay events. For each event, spike rasters (ordered by the location of each neuron’s place field) and the Bayesian decoded trajectory are shown. “+” (“−”) label corresponds to significant (insignificant) events. (left) Both methods can fail to label events that appear to be sequential as replay and (right) label events replay that appear non-sequential. **b.** We recruited human scorers to visually inspect Bayesian decoded spike trains and identify putative sequential replay events. Using their identifications as labels, we can define an ROC curve for both Bayesian and HMM model-congruence which shows how detection performance changes as the significance threshold is varied. (inset) Human scorers identify 24% of PBEs as replay. Setting thresholds to match this value results in agreement of 70% between Bayesian and HMM model-congruence. **c.** Using the same thresholds, we find ≈ 70% agreement between algorithmic and human replay identification. (All comparison matrices, *p* < 0.001, Fisher’s exact test two-tailed.) Figure 5-Figure supplement 1. Human scoring of PBEs and session quality.

We marked each event as a “true” community replay if it was identified by a majority of scorers (six individuals scored *n* = 1883 events, two individuals scored a subset of *n* = 1423 events, individual scores are shown in *Figure 5-Figure Supplement 1*). We calculated an receiver operating characteristic (ROC) curve which compared the rate of true positive and false positive detections as the significance thresholds for Bayesian and model-congruence approaches were varied (***Figure 5***b). A perfect detector would have an area under the curve (AUC) of unity. We did not find a significant difference between the AUCs of Bayesian decoding and model-congruence (*p* = 0.14, bootstrap, see Methods). If we select thresholds such that our algorithms yield a similar fraction of significant vs. total events as the 24% denoted by our human scorers, we find that both Bayesian and model-congruence yield agreement of ≈ 70% labeled events with each other and with human scorers (***Figure 5***c).

Thus, congruence with an HMM trained only on PBEs appears to work as reliably as Bayesian decoding in detecting sequential reactivation of linear track behaviors. However, when we examined individual sessions, we noticed that our performance was quite variable. Given that our models are learned only from PBEs, we reasoned that the statistics or structure of the PBEs within each session might yield models which varied in quality. We created a model quality metric by comparing cross-validated learning statistics to models which were learned from shuffled events (see Methods). We found that the performance of model-congruence detection was tied to model quality (*R*^2^ = 0.26, *F* = 4.591, *n* = 18 sessions, *Figure 5-Figure Supplement 1*). Model quality, in turn, was highly correlated with the number of PBEs during the session (*R*^2^ = 0.92, *F* = 191.6, *n* = 18 sessions, *Figure 5-Figure Supplement 1*). Not surprisingly, there wasn’t a relationship between model quality or the number of PBEs and Bayesian decoding performance because the place field model is learned from ensemble neural activity during running. Thus, we find an intriguing contrast—when there is an abundance of PBEs (indicating novelty, learning, hippocampally-dependent planning, etc.), even in the absence of repeated experience, replay detection based on PBE activity is highly effective. Conversely, when there are few PBEs (i.e., scenarios in which PBEs are uncorrelated with cognitive function), but an abundance of repeated behavioral trials, Bayesian decoding of these limited events proves more effective.

### Modeling internally generated activity during open field behavior

The linear track environment represents a highly-constrained behavior. We therefore asked whether the hidden Markov model approach would generalize to more complex environments and behavioral tasks. ***Pfeiffer and Foster*** (***2013***, 2015) had previously recorded activity in CA1 in rats as they explored in a 2 m × 2 m open field arena for liquid reward. Briefly, animals were trained to discover which one of 36 liquid reward wells would be the “home” well on a given day. They then were required to alternate between searching for a randomly rewarded well and returning to the home well. Using the place cell map in this task and Bayesian decoding, many PBEs were decoded to trajectories through two-dimensional space that were predictive of behavior and shaped by reward. Using this same dataset, we trained an HMM on neural activity during PBEs in the open field. Here, we used the same PBEs detected previously (*Pfeiffer and Foster, 2013*) which occupied an average of 2.53±0.42% of the periods during which animals were behaving (77.91±21.16 s out of 3064.86±540.26 s). Given the large number of units available in this dataset and the increased behavioral variability in the open field environment compared to the linear track, we chose to estimate HMMs with 50 latent states. The transition matrix and observation model from a sample session are shown in ***Figure 6***a,b. Despite the complex and varied trajectories displayed by animals, the HMM captured sequential dynamics in PBE activity—as in the 1D case, when we compared learned models against both actual and surrogate test data and found that the model likelihood was significantly greater for real data (*p* < 0.001, Wilcoxon signed-rank test).

**Figure 6.**
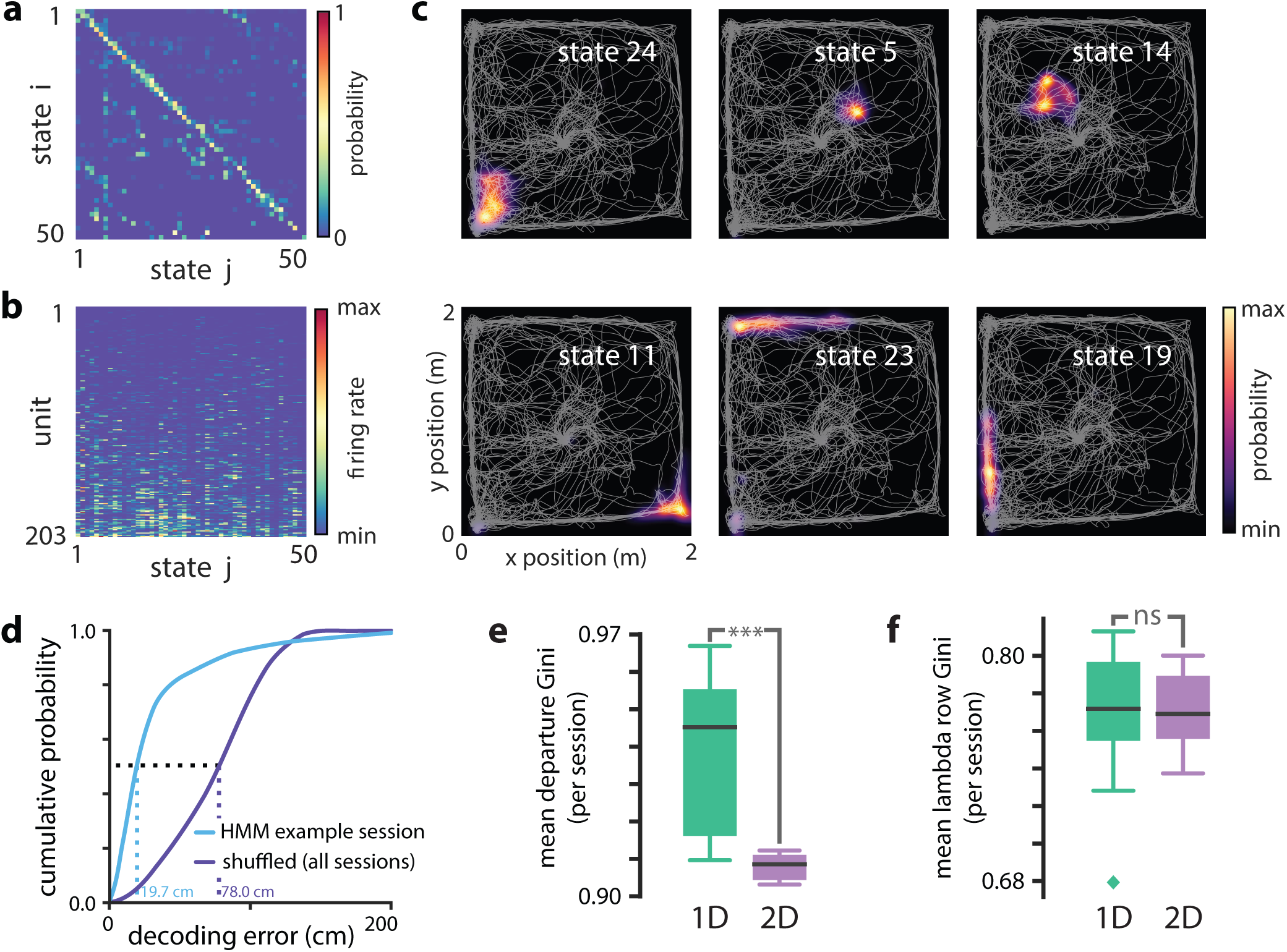
Modeling PBEs in open field. **a.** The transition matrix estimated from activity detected during PBEs in an example session in the open field. **b.** The corresponding observation model (203 neurons) shows sparsity similar to the linear track. **c.** Example latent state place fields show spatially-limited elevated activity in two dimensions. **d.** For an example session, position decoding through the latent space gives significantly better accuracy than decoding using the shuffled latent state place fields. **e.** Comparing the mean departure Gini coefficient per session between the linear track and open field reveals that, as expected, the open field is significantly *less sparse* (*p* < 0.001), since the environment is less constrained. However, **f.** shows that there is no significant difference between the mean observation Gini coefficient (across states) between the linear track and the open field. Figure 6-Figure supplement 1. Open field PBE model states typically only transition to a few other states. Figure 6-Figure supplement 2. Each neuron is active in only a few model states in the open field. Figure 6-Figure supplement 3. lsPFs and position decoding in an open field. Figure 6-Figure supplement 4. Examples of open field PBEs.

In the case of the linear track, we linked sparsity of the transition matrix to the sequential nature of behaviors in that environment. An unconstrained, two-dimensional environment permits a much richer repertoire of behavioral trajectories. However, behavior is still constrained by the structure of space—arbitrary teleportation from one location to another is impossible. We found that learning from PBEs in the open field yielded transition matrices (***Figure 6***a) that were significantly more sparse than models learned from shuffled data (*p* < 0.05, Wilcoxon signed-rank test, *n* = 8 sessions, *Figure 5-Figure Supplement 1*). However, consistent with increased freedom of potential behaviors, when we compared the sparsity of models learned from open field PBEs with 50-state models learned from PBEs in linear tracks, the open field transition matrices were less sparse (*p* < 0.001, Mann-Whitney *U* test comparing 8 and 18 sessions, *Figure 4-Figure Supplement 1*). Likewise, when we examined the observation model for the open field, we found that the activity across states for individual neurons was significantly more sparse than in models learned from shuffled data (*p* < 0.05, Wilcoxon signed-rank test, *n* = 8 sessions, *Figure 6-Figure Supplement 1*). Consistent with the idea that the neural ensemble in the dorsal hippocampus should be consistent, the sparsity of linear track and open field observation models were not significantly different (*p* = 0.44, Mann-Whitney *U* test).

Can an HMM trained on PBEs indeed capture spatial information about a 2D environment? We used the PBE-trained model to decode run data to calculate lsPFs, as in the linear track case. We found that the latent states corresponded with specific locations in the open field (***Figure 6***c). Moreover, we were able to decode animals’ movements with significantly greater than chance accuracy by converting decoded latent states to positions using the lsPF (*p* < 0.001, ***Figure 6***d). Finally, we examined model-congruency for PBEs detected in the open field. Previously, it was reported that 27.3% (815 of 2980, *n* = 8 sessions) were identified as “trajectory events” (*Pfeiffer and Foster, 2015*). We chose a significance threshold to match this fraction (*Figure 6-Figure Supplement 3*) and found that there was significant overlap between the events detected through Bayesian and model-congruence techniques (*p* < 0.01, Fisher’s exact test). These events were significantly overlapped with replay events detected using traditional Bayesian decoding (*Figure 6-Figure Supplement 3*). Thus, an HMM of the activity during population bursts captures the structure of neural activity in two dimensional environments during complex tasks and can be used to decode events consistent with trajectories through that environment.

## Discussion

Increasing lines of evidence point to the importance of hippocampal ensemble activity during PBEs in guiding on-going behavior and active learning. Despite being the strongest output patterns of the hippocampus, however, this activity has been assumed to be interpretable only in the context of other theta-associated place cell activity expressed during behavior. Our findings demonstrate that over the course of a behavioral session, ensemble activity during PBEs alone is sufficient to form a model which captures the spatial relationships within an environment. This suggests that areas downstream of the hippocampus might be able to make use solely of PBE activity to form models of external space. In an extreme view, place cell activity might merely subserve the internal mechanisms in the hippocampus which generate PBE sequences. To the extent that animals might wish to use the spatial code obtained from PBEs to identify their current location, we show that this can be done after translating ensemble activity into the latent states of the model.

When we examined the transition matrices we learned from PBEs, we found that they were marked by significant sparsity. This sparsity results from the sequential patterns generated during PBEs. Latent variable models have previously been used to analyze the structure of hippocampal place cell activity (*Chen et al., 2012*, *2014; Dabaghian et al., 2014*). In these studies, the learned transition matrices were mapped to undirected graphs which could be analyzed using topological measures. It is intriguing that similar structure is apparent in PBE activity. For example, we observed that transition matrices learned from PBEs associated with linear track behavior were significantly more sparse than those learned from the open field, which we hypothesize is a consequence of the greater freedom of behavior in the latter (a topological difference). Whether hippocampal PBE activity must always be sequential, i.e., evolve through a sparsely-connected latent space, is an open an interesting question, as are differences between the latent state space dynamics learned during PBEs and those learned from place cell activity.

### Graded, Non-binary Replay Detection

Remarkably, evaluating the congruence or likelihood of test data against our HMM provided a highly novel method to detect events that are consistent with replay, without a need to access the “play” itself. In the process of evaluating the potential of HMMs for detecting replay, we developed an approach to compare different replay-detection strategies. Our results highlight how the data does not readily admit to a strict separation between “replay” and “non-replay” events. While it is possible that with additional shuffles or other restrictions (*Silva et al., 2015*), automated performance might be rendered closer to human-labeling, even human scorers had variation in their opinions. This calls into doubt judgments of memory-related functions which build on a binary distinction between replay and non-replay sequences. Model congruence, either as a raw statistical likelihood or weighted against a shuffle distribution, seems to be a very reasonable metric to associate with individual PBEs. Moreover, evaluating congruence with an HMM does not require access to repeated behavioral sequences, which may be unfeasible under widely-used single- or few-trial learning paradigms or when the events involve replay of a remote internalized environment. Given these benefits, along with computational efficiency, we would suggest that future analyses of the downstream impact of hippocampal reactivation regress effects against this measure rather than assuming a binary distinction.

### Learning, Model Congruence and Replay Quality

Not surprisingly, the rate of PBEs had a large effect on our ability to measure model congruence. Interestingly, it has been noted that the density of PBEs is higher during early exposure to a novel environment (*Cheng and Frank, 2011; Frank et al., 2004; Kemere et al., 2013; Kudrimoti et al., 1999*). This might suggest that for the animal, PBE activity would be an important source for generating models of the world when the animal is actively learning about the environment. If as hypothesized, replay is a form of rehearsal signal generated by the hippocampus to train neocortical modules (*McClelland et al., 1995*), then indeed the brain’s internal machinery may also be evaluating whether a given sequential PBE pattern is congruent and consistent with previously observed PBEs. In later sessions, as animals have been repeatedly exposed to the same environments, downstream regions will have already witnessed many PBEs from which to estimate the structure of the world. Overall, our approach provides a novel viewpoint from the perspective of hippocampal PBEs. An interesting future line of inquiry would be to assess the extent to which a model built on PBEs during first experience of a novel environment is slower or faster to converge to the final spatial map than models built on theta-associated place activity.

### Application to Non-spatial Behaviors

We have analyzed data gathered in experiments in which rats carried out simple spatial navigation tasks. Thus, it is to some extent not surprising that when we decoded ensemble activity during behavior we found that spatial position was strongly associated with the latent states. However, our approach could easily be applied to behaviors in which tasks involve variables other than position as well as tasks in which spatial information has no role. While following our approach for calculating latent-state place fields might be challenging apply in these cases, assessing replay as model-congruence would be simple. This points to future studies in which the role of sequential but non-spatial hippocampal replay might be assessed.

### Future possibilities

It has been previously observed that the rate of hippocampal reactivations in PBEs during awake behavior is much higher than during sleep (*Grosmark and Buzsáki, 2016; Karlsson and Frank, 2008*), but the reasons for this are not well understood. One hypothesis is that many sleep PBEs contain the reactivation of contexts other than those measured during a behavioral experiment. Another hypothesis is that sleep activity involves remodeling of dynamic network architectures (*Buhry et al., 2011; Tononi and Cirelli, 2014*). Our approach has the potential to illuminate some sources of variability during sleep. With sufficient data, our HMM approach should be able to learn disjoint sets of latent states (or “sub-models”) which would capture these separate contexts and allow us to test this possibility. Note that we would also expect multiple sub-models in awake PBEs in data in which remote awake replay has previously been observed (*Karlsson and Frank, 2009*). If instead of such sub-models, sleep PBEs yielded parameters which were markedly different (e.g., less sparse) than those learned from awake PBEs, that might support the network remodeling function of sleep. In the latter case, we might imagine that only a small subset of sleep PBEs—corresponding to learning-related replay—would be congruent with a model learned from awake PBE data.

### Conclusions

We have demonstrated a new analytical framework for studying hippocampal ensemble activity which enables primacy of PBEs in model formation. We use an unsupervised learning technique commonly used in the machine learning field to study sequential patterns, the hidden Markov model. This contrasts with existing approaches in which the model—estimated place fields for the ensemble—is formed using the theta-associated place cell activity. We find that our PBE-frst approach results in a model which still captures the spatial structure of the behavioral tasks we studied. Additionally, we demonstrate that we can use model-congruence as a tool for assessing whether individual PBEs contain hippocampal replay. This analytical approach bears much promise for illuminating a number of areas in which traditional analyses have faltered.

## Materials and Methods

### Experiment paradigm/Neural data recording

We analyzed neural activity recorded from the hippocampus of rats during periods in which they performed behavioral tasks for food reward in two paradigms. First, we considered data from animals running back and forth in a linear track 150 or 200 cm long. As previously reported (*Diba and Buzsáki, 2007*), we recorded neural activity using chronically-implanted silicon probes to acquire the activity of hippocampal CA1/CA3 neurons. From these experiments, we chose sessions during which we observed at least 20 place cells during active place-field exploration, and at least 30 PBEs (see below). Place cells were identified as pyramidal cells which had (i) a minimum peak firing rate of of 2 Hz, (ii) a maximum mean firing rate of 5 Hz, and (iii) a peak-to-mean firing rate ratio of at least 3, all estimated exclusively during periods of run (as defined before, that is, when the animal was running > 10 cm/s). This selection yielded *n* = 18 session with 41–203 neurons (36–186 pyramidal cells). All procedures were approved by the Institutional Animal Care and Use Committee of Rutgers University and followed US National Institutes of Health animal use guidelines.

Second, we considered data recorded using tetrodes to record a large number (101–242) of putative pyramidal neurons in area CA1 during two sessions each in four rats. Briefly, as was previously reported (*Pfeiffer and Foster, 2013*), rats explored an arena in which there were 36 reward sites. In each session, one site was designated as “home”. During a session, rats would repeatedly alternate between retrieving a random reward site in one of the remaining 35 locations and retrieving a reward at the home location. All procedures were approved by the Johns Hopkins University Animal Care and Use Committee and followed US National Institutes of Health animal use guidelines.

### Population burst events

To identify PBEs in the linear track data, a spike density function (SDF) was calculated by counting the total number of spikes across all recorded single and multi-units in non-overlapping 1 ms time bins. The SDF was then smoothed using a Gaussian kernel (20 ms standard deviation, 60 ms half-width). Candidate events were identified as time windows with a peak SDF of at least three standard deviations above the mean calculated over all the session. The boundaries of each event were set to time points of crossing the mean, preceding and following the peak. Events during which animals were moving (average movement speed of > 5 cm/s) were excluded from all further analyses to prevent possible theta sequences from biasing our results. The activity of putative interneurons (mean firing rate when moving > 10 Hz) was excluded. For analysis, we then binned each PBE into 20 ms (non-sliding) time bins. Finally, events with duration less than four time bins or with fewer than 4 active pyramidal cells. For the open field data, we used the previously reported criteria for identifying PBEs prior to binning (10 ms standard deviation kernel, minimum of 10% of units active, duration between 50 ms and 2000 ms).

### Hidden Markov model of PBE activity

We trained HMMs on the PBEs. In an HMM, an unobserved discrete latent state *q_t_* evolves through time according to a first order Markov process. The temporal evolution of the latent state is described by the *M* × *M* matrix **A**, whose elements {*a_ij_*} signify the probability after each time bin of transitioning from state *i* to state *j*, *a_ij_* = Pr (q*_t_*_+1_ = *j*|*q****_t_*** = *i*). The number of states, *M*, is a specified hyperparameter. We found that our results were insensitive to the value of *M* through a wide range of values from 20 to 100 (*Figure 3-Figure Supplement 1*). During each time bin of an event, the identity of the latent state influences what is observed via a state-dependent probability distribution. We modeled the *N*-dimensional vector of binned spiking from our ensemble of *N* neurons at time *t*, *O_t_*, as a Poisson process. Specifically, for each state, *i*, we model neuron *n* as independently firing according to a Poisson process with rate *λ_ni_*.

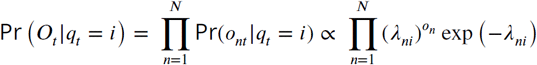

where *o_nt_* is the number of spikes observed from neuron *n* at time *t*. The final parameter which specifies our model is the probability. distribution of the initial state for a given event, *π_i_* = *Pr*(*q*_1_ = *i*). Thus, our model is specified by parameters *θ* = {**A**, **Λ**, ***π***}, where **Λ**= {*λ_ni_S*} is an *N* × *M* matrix and ***π*** = {*π_i_*} is an *N*-dimensional vector.

To learn model parameters, we follow the well-known iterative EM procedure (*Rabiner, 1989*), treating each training PBE as an observation sequence. In order to regularize the model, we impose a minimum firing rate for each neuron of 0.001 (0.05 Hz) during the M-step of EM. For a given PBE (i.e., observation sequence) with *K* bins, we use the “forward-backward algorithm” (*Rabiner, 1989*) to calculate both the probability distribution of the latent state for each time bin, Pr(*q_t_*|*O*_1_, …, *O_t_*, …, *O_K_*), and the “score”, or likelihood of the complete sequence, Pr(*O*_1_, …, *O_K_*). We define the model likelihood of an HMM as the product of the scores of test sequences using five-fold cross validation. To initially evaluate model learning, we compared model likelihoods calculated using real and shuffled test data. Models which have learned to properly represent the data should show significant increases.

### Ordering states for visualization

For visualization, we wanted to order the states to maximize the super diagonal of the transition matrix. We used a greedy approach which typically yields this solution. We started by assigning the first index to the state with the highest initial probability and added states based on the most probable state transitions. The undirected connectivity graphs were then generated from this transition matrix, averaging the strength of reciprocal connections, *a_ij_* and *a_ji_*.

### Surrogate datasets and shuffle methods

We obtained three types of surrogate datasets by shuffling the neural activity during PBEs in distinct ways. 1) Temporal shuffle: within each event, the binned spiking activity was circularly permuted across time for each unit, independently of the other units. This goal of this shuffle is to disrupt unit co-activation, while maintaining the temporal dynamics for each unit. 2) Time-swap shuffle: within each event, the order of the binned columns of neural activity was randomly permuted across time, keeping the unit co-activation intact. The goal of this shuffle is to change the temporal dynamics of ensemble activity. 3) Poisson surrogate “shuffle”: we estimated each unit’s mean firing rate across all PBEs, and then produced surrogate PBEs from independent Poisson simulations according to each unit’s mean firing rat.

### Calculating sparsity and connectivity of the model parameters

Sparsity of the transitions from individual states (departure sparsity) was measured by calculating the Gini coefficient of corresponding rows of the transition matrix (*Hurley and Rickard, 2009*). For analyses of PBE models from linear tracks, we computed the mean sparsity across states for each of the 250 surrogate datasets, and these means were used to generate the box plots of ***Figure 2***c. Note that for the actual data, we generate a distribution by randomly initializing the model 250 times and calculating the mean sparsity for each initialization. For analyses of models learned from PBEs in open fields (and the linear track comparison with 50 states), we created 50 surrogates/random initializations (*Figure 6-Figure Supplement 1*). To compare across sessions, we calculated the mean sparsity by averaging over all 250 surrogate datasets to obtain a single mean sparsity per session, so that *n* = 18 per-session means were used to create the box-plots of ***Figure 2***e.

Firing rates for different units can be highly variable. Thus, when evaluating the sparsity of the observation matrix, we measured the extent to which individual units were active in only a few states by calculating the Gini coefficients of the rows of the observation matrix. As with transitions, we calculated mean sparsity across units for each surrogate datasets to generate ***Figure 2***d (open field, *Figure 6-Figure Supplement 2*), and we then averaged over the surrogate datasets to obtain a per-session average used in ***Figure 2***f.

### Model connectivity and sequences

To measure the degree of sequential connectivity within the graph corresponding to the transition matrix—with nodes and edges representing the states and transitions, respectively—we developed an algorithm for measuring the length of the longest path that can be taken by a random walk on the graph. To do this, we made an adjacency matrix for a corresponding unweighted directed graph by binarizing the transition matrix using a threshold of 0.2 on the transition probabilities. Starting from each node, then, the algorithm finds the longest path that ends to either a visited or a terminal node (a node without any outgoing edges). In case of comparison between models trained on actual data and each of the surrogate datasets, we adjusted the thresholds in a way to match the average degree—the average number of edges per node—between the models, ruling out possible effects due to any differences in the total number of graph edges. We carried out the analysis on the same set of models previously generated for analyzing sparsity. To compare across sessions, we calculated the median maximum path length for each session (*n* = 18) and used the per-session medians to generate box plots of *Figure 2-Figure Supplement 3*c.

### Latent State Place Fields

To calculate the latent state place fields, we first identified bouts of running by identifying periods when animals were running (speed > 10 cm/s). We then binned the spiking during each of these bouts in 100 ms bins. Using the forward-backward algorithm and the HMM model parameters learned from PBEs, we decoded each bout into a sequence of latent state probability distributions, Pr(*q_t_*|*O_t_*). After binning the track positions corresponding to each time bin, we then found the average state distribution for each position bin, *x_p_*, and normalized to yield a distribution for each state, Pr(*x_p_*|*q_t_* = *i*).

### Decoding position from latent state trajectories

In order to decode the animal’s position during bouts of running using decoded latent state trajectories, we use the lsPFs. With five-fold cross validation, we estimated lsPFs using a training data set, then use the PBE model to decode latent state distributions from ensemble neural activity in the test data. The product of lsPFs and decoded latent state distribution at time *t* is the joint distribution Pr(*x_p_*, *q_t_*|*O_t_*). We decode position as the mean of the marginal distribution Pr(*x_p_*|*O_t_*).

### Bayesian Replay Detection

In order to detect replay in our 1D data, we used a Bayesian decoding approach (*Kloosterman, 2012*). For each 20 ms time bin *t* within a PBE, given a vector comprised of spike counts from *N* units, *O_t_* = (*o*_1_*_t_ o*_2_*_t_* … *o_Nt_*) in that bin, the posterior probability distribution over the binned track positions was calculated using the Bayes’ rule:

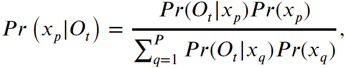

where *x_p_* is the center of *p*-th linearized position bin (of *P* total). We assume Poisson firing, thus the prior probability, Pr(*O_t_*|*x_p_*,), for the firing of each unit *n* is equal to

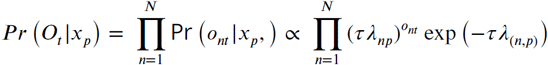

where *τ* is the duration of time bin (100 ms during estimation, 20 ms during decoding), and *λ_np_* characterizes the mean firing rate of the *n*-th unit in the *p*-th position bin. We assumed a uniform prior distribution *Pr*(*x_p_*) over the position bins.

For each PBE, the estimated posterior probability distribution was used to detect replay as follows. Many (35,000) lines with different slopes and intercepts were sampled randomly following the approach in (*Kloosterman, 2012*). The Bayesian replay score for a given event was the maximum score obtained from all candidate lines, where the score for a particular line was defined as the mean probability mass under the line, within some bandwidth (a halfwidth of 3 cm was used here). For time bins during which the sampled line fell outside of the extent of the track, the median probability mass of the corresponding time bin was used, and for time bins during which no spikes were observed, we used the median probability mass across all on-track time bins. To evaluate the significance of this score, for each event we generated 5,000 surrogates of the posterior probability distribution by cycling the columns (i.e., for each time bin, circularly permuting the distribution over positions by a random amount). For each of these surrogates a replay score was calculated. The Monte Carlo p-value for each event was calculated as the number of shuffled events with replay scores higher than the raw data. The threshold for significance was varied as described in the text. For the open field, we used previously reported criteria (*Pfeiffer and Foster, 2013*) to identify replay events from PBEs.

### Replay detection via PBE model congruence

To identify replay as model congruence, we used the HMM sequence score as defined above. Using five-fold cross validation, the parameters of a HMM were learned from training PBEs. The sequence score was then calculated for each event in the test data. To evaluate the significance of this score, for each event we generated 5,000 surrogate scores using a computationally-efficient scheme. Specifically, for each surrogate, we randomly shuffle the rows of the transition matrix, excepting the diagonal. By maintaining the diagonal and leaving the observation model unchanged, this shuffle specifically selects against PBEs in which the latent states do not evolve in temporal sequences. The Monte Carlo *p*-value for each event was calculated as the number of shuffled events with HMM sequence scores higher than the raw data. The threshold for significance was varied as described in the text.

### Human scoring and detection comparison

We organized a group of human scorers to use visual inspection to evaluate whether individual PBEs should be described as replay. More specifically, scorers were only presented with Bayesian decoded probability distributions such as those in ***Figure 4***a, but without access to the spike raster or any additional information. We built an automatic presentation system which would display each event in random order, and record one of six possible scores: “excellent” (highly sequential with no jumps and covering most of the track), “good” (highly sequential with few or no jumps), “flat” (decoded position stayed mostly in the same place, i.e. no temporal dynamics), “uncertain” (some semblance of structure, but not enough to fall into any of the previous categories) or “noise” (no apparent structure, or nonsensical trajectories such as teleportation). For analysis, an event was designated as replay if it was labeled as “excellent” or “good” by a majority of scorers.

To calculate an ROC curve for replay detection algorithms, we used our shuffle statistics for each event to create a vector which related the significance threshold (e.g., 99%) to the label applied by the algorithm’s (i.e., significant replay or not). Then, as a function of threshold, the sensitivity (fraction of true positives identified) and selectivity (fraction of true negatives identified) was averaged over events to yield an ROC curve. To evaluate whether the AUC differed between Bayesian and model-congruence techniques we used a bootstrap approach. To generate a null hypothesis, we combined the event/threshold vectors from both groups, and then sampled with replacement two random groups (A and B) from the pooled data with replacement. The AUCs for these two random groups of events were measured, and a distribution for the difference between the randomly chosen AUCs was calculated. The two-sided *p*-value we report is the fraction of differences in random AUCs which are more extreme than the actual difference.

### HMM model quality across sessions

In order to understand the extent to which an HMM trained on PBEs from a given session contained sequentially-structured temporal dynamics, we learned models on surrogate cross-validated data sets derived by shuffling the actual data using the time-swap shuffle. In particular, using five-fold cross validation, we calculated the likelihood of each event in the session (that is, for each fold, we learn an HMM on the training set, and score each event in the validation / test set using this HMM). Subsequently, we repeat this same process, but on time-swap shuffled data. We repeated this process *n* = 200 times (each time doing five-fold cross validation, and learning an HMM per fold), so that we end up with a distribution of event scores coming from shuffled test data evaluated in shuffled-optimized models, and our actual data event scores, evaluated in our actual-data-optimized model. Then, we define the session quality as the average z-score of the actual data events, compared to the shuffled data distribution.

## Acknowledgments

This work was supported by the NSF (IOS-1550994 and CBET-1351692) the Human Frontiers Science Program, the NIMH (R01MH109170 and R01MH085823), the Alfred P. Sloan Foundation, the Brain and Behavior Research Foundation, the McKnight Endowment Fund for Neuroscience, and the University of Texas System. Furthermore, we are grateful for the volunteer human scorers, Sibo Gao, Ariel Feldman, Eric Lewis, Joshua Chu, Kathleen Hu, Shayok Dutta, and James Webb, who patiently viewed hundreds of sequences.

---

**Figure 1-Figure supplement 1.**
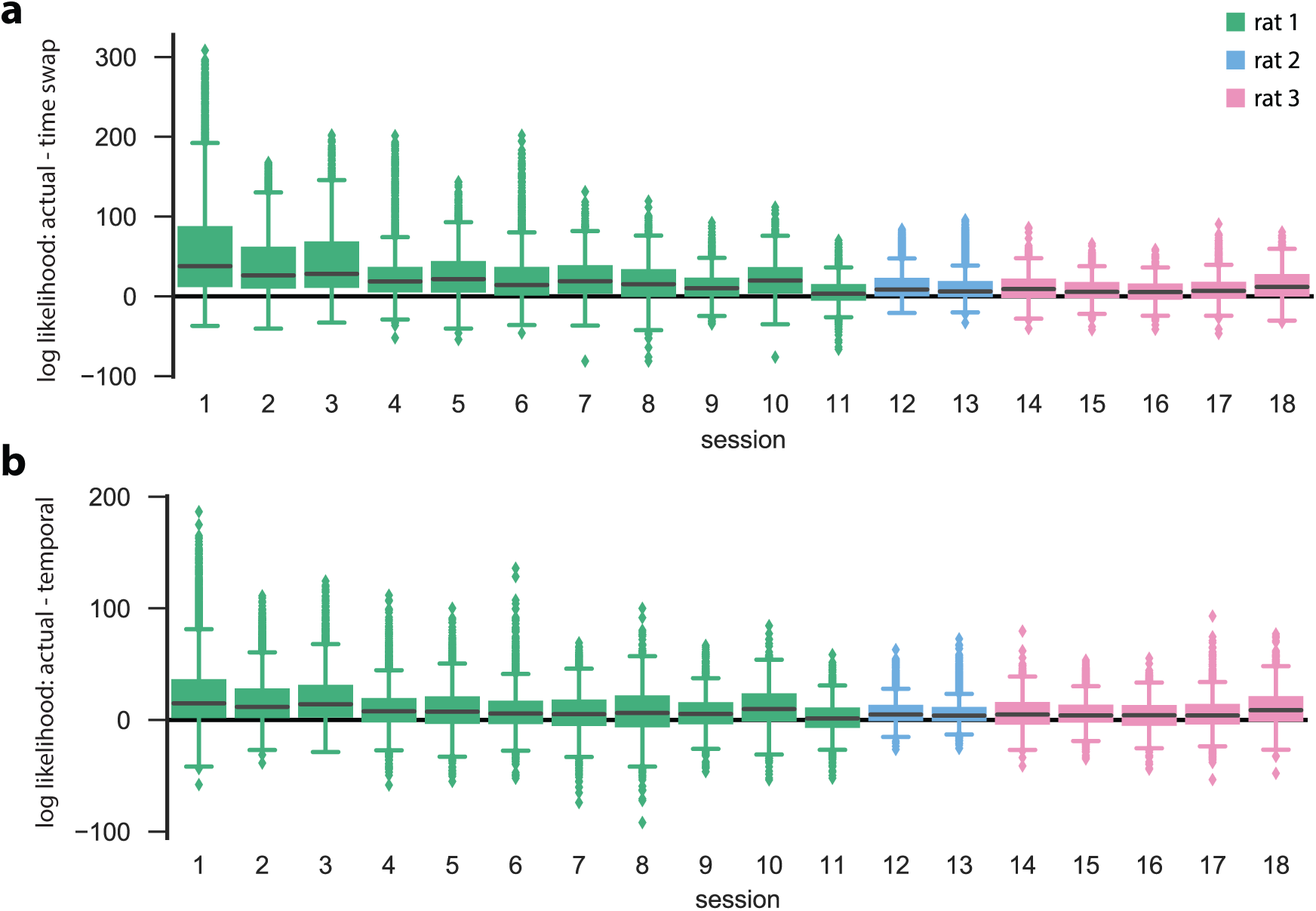
Actual cross-validated test data and surrogate test data evaluated in actual-data-optimized HMMs for all 18 linear track sessions. For each session, we performed five-fold cross validation to score the validation (=test) set in an HMM that was learned on the corresponding training set. In addition, two surrogate datasets of the validation data (obtained by either temporal shuffle or time swap shuffle) were scored in the same HMM as the actual validation data. *k* = 50 shuffles of each event and of each type were performed. **a.** Difference between the data log likelihoods of actual and time swap surrogate test events, evaluated in the actual train-data-optimized models. **b.** Same as in **a.**, except that the differences between the actual data and the temporal surrogates are shown. For each of the *n* = 18 sessions, the actual test data had a significantly higher likelihood than either of the shuffled counterparts (*p* < 0.001, Wilcoxon signed-rank test). Sessions are arranged first by animal, and then by number of PBEs, in decreasing order.

**Figure 2-Figure supplement 1.**
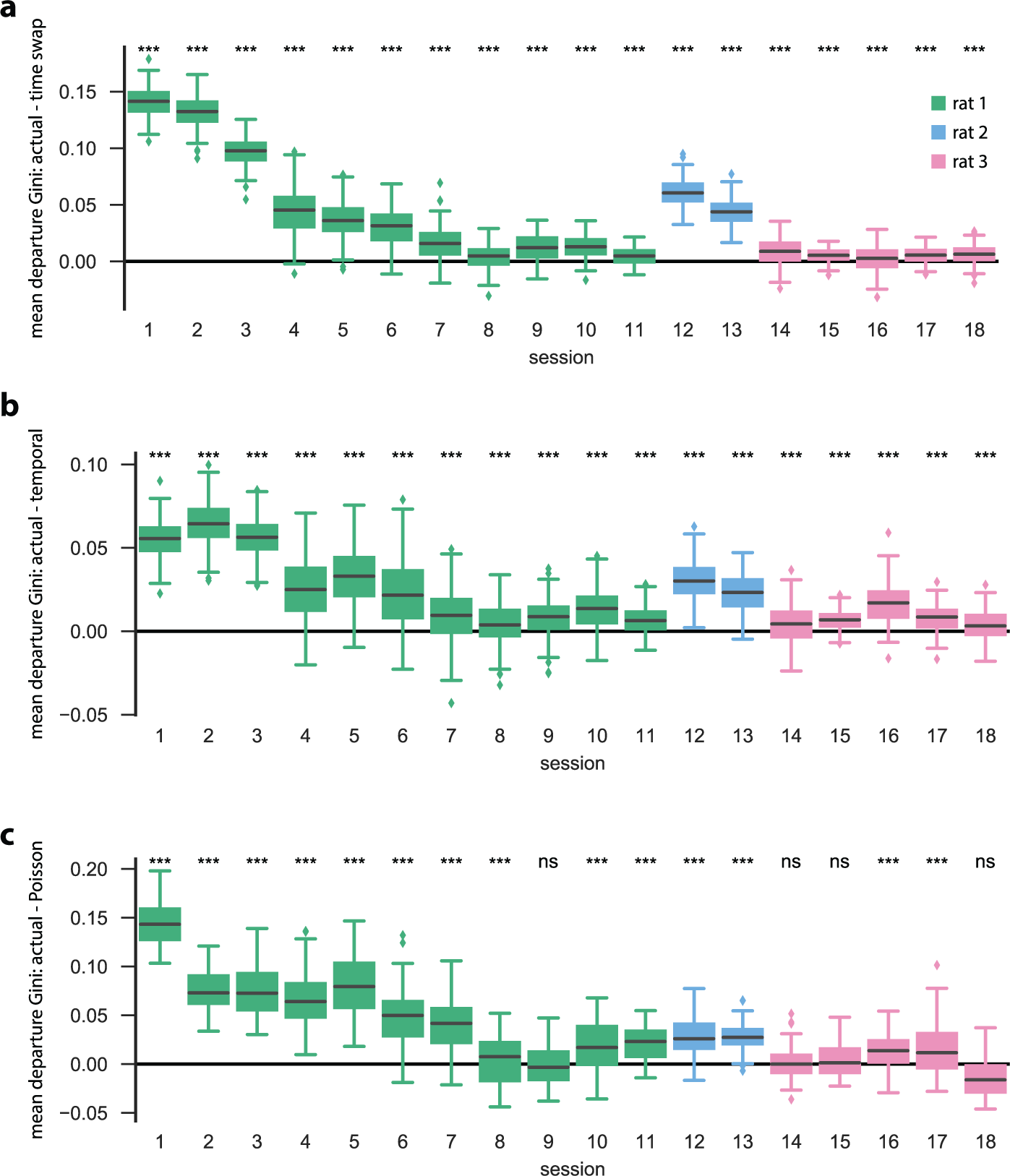
We trained HMMs on neural activity during PBEs (in 20 ms bins), and asked how sparse the resulting state transitions were. In particular, we calculated the Gini coefficient for each row of our state transition matrix, so that the Gini coefficient for a particular row reflects the sparsity of state transitions from that state (row) to all other states (so-called “departure sparsity”). A high (close to one) Gini coefficient implies that the state is likely to only transition to a few other states, whereas a low (close to zero) Gini coefficient implies that the state is likely to transition to many other states. For each transition matrix, we computed the mean departure sparsity for *N* = 250 initializations, and for *N* = 250 shuffled counterparts for each of the surrogate data sets (**a.** time swap shuffle, **b.** temporal shuffle, **c.** Poisson surrogate), and in each case we show the difference between the actual test data, and the surrogate test data. The actual data are significantly more sparse than both the temporal and time swap surrogates for all sessions (*p* < 0.001, Mann-Whitney *U* test) and significantly more sparse than the Poisson surrogate for 14 of the 18 sessions (*p* < 0.001, Mann-Whitney *U* test).

**Figure 2-Figure supplement 2.**
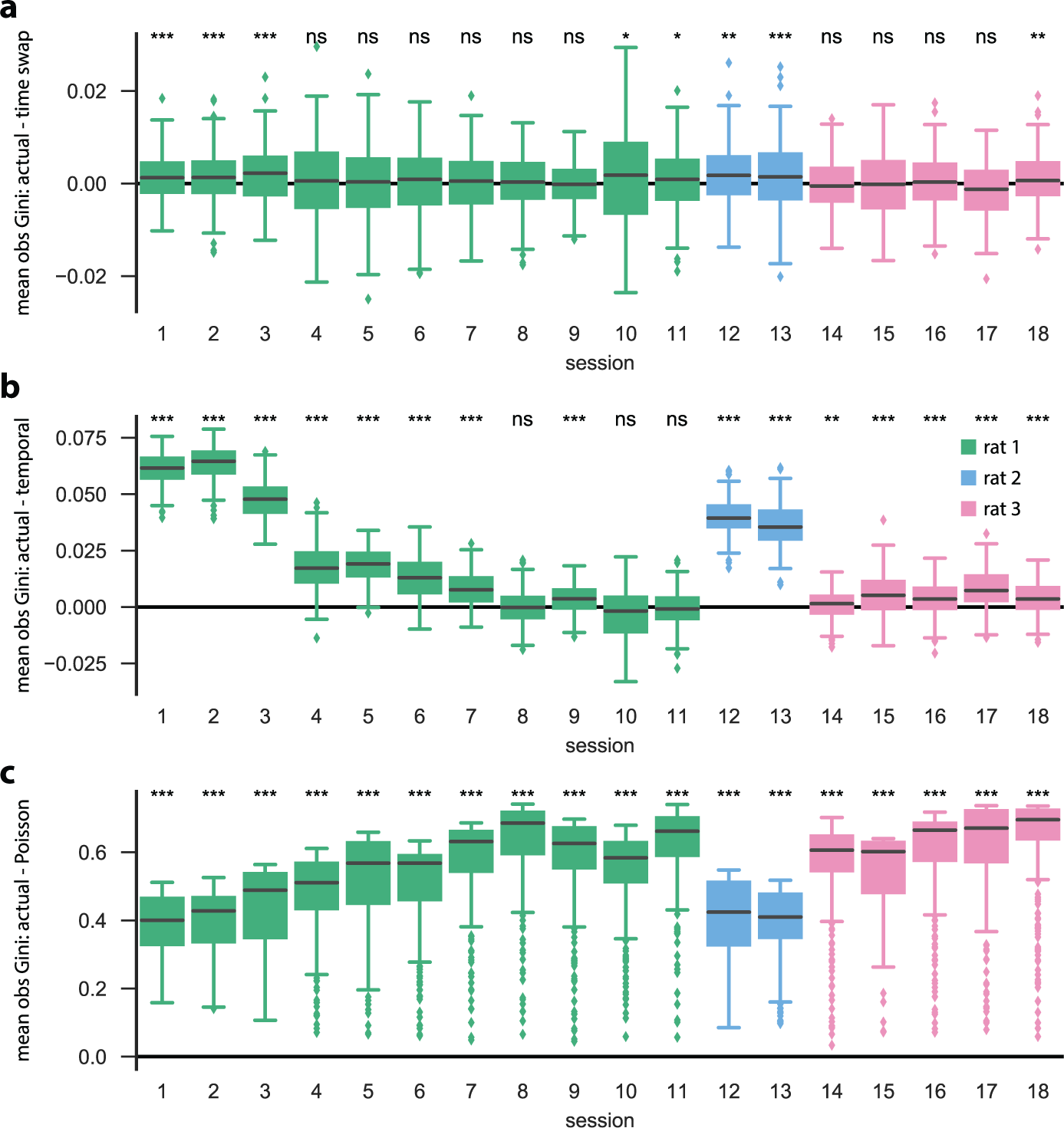
Using the same PBE models and surrogate datasets (N=250 shuffles each) as in *Figure 2-Figure Supplement 1*, we investigated the sparse participation of neurons/units in our models by calculating the Gini coefficient of each row (that is, for each unit) of the observation matrix. A high Gini coefficient implies that the unit is active in only a small number of states, whereas a low Gini coefficient implies that the unit is active in many states. For each initialization / shuffle, we calculate the mean Gini coefficient over all units, and the differences between those obtained using actual data and those obtained using surrogate data are shown: differences between actual and **a.** time swap, **b.** temporal, and **c.** Poisson surrogates. We find that the actual data are significantly more sparse than the temporal and Poisson surrogates for most of the sessions (p < 0.001, Mann-Whitney *U* test), but that for many (10 out of 18) sessions, there is no significant difference between the mean row-wise observation sparsity of the actual data compared to the time swap surrogate. This is an expected result, since the time swap shuffle leaves the observation matrix largely unchanged.

**Figure 2-Figure supplement 3.**
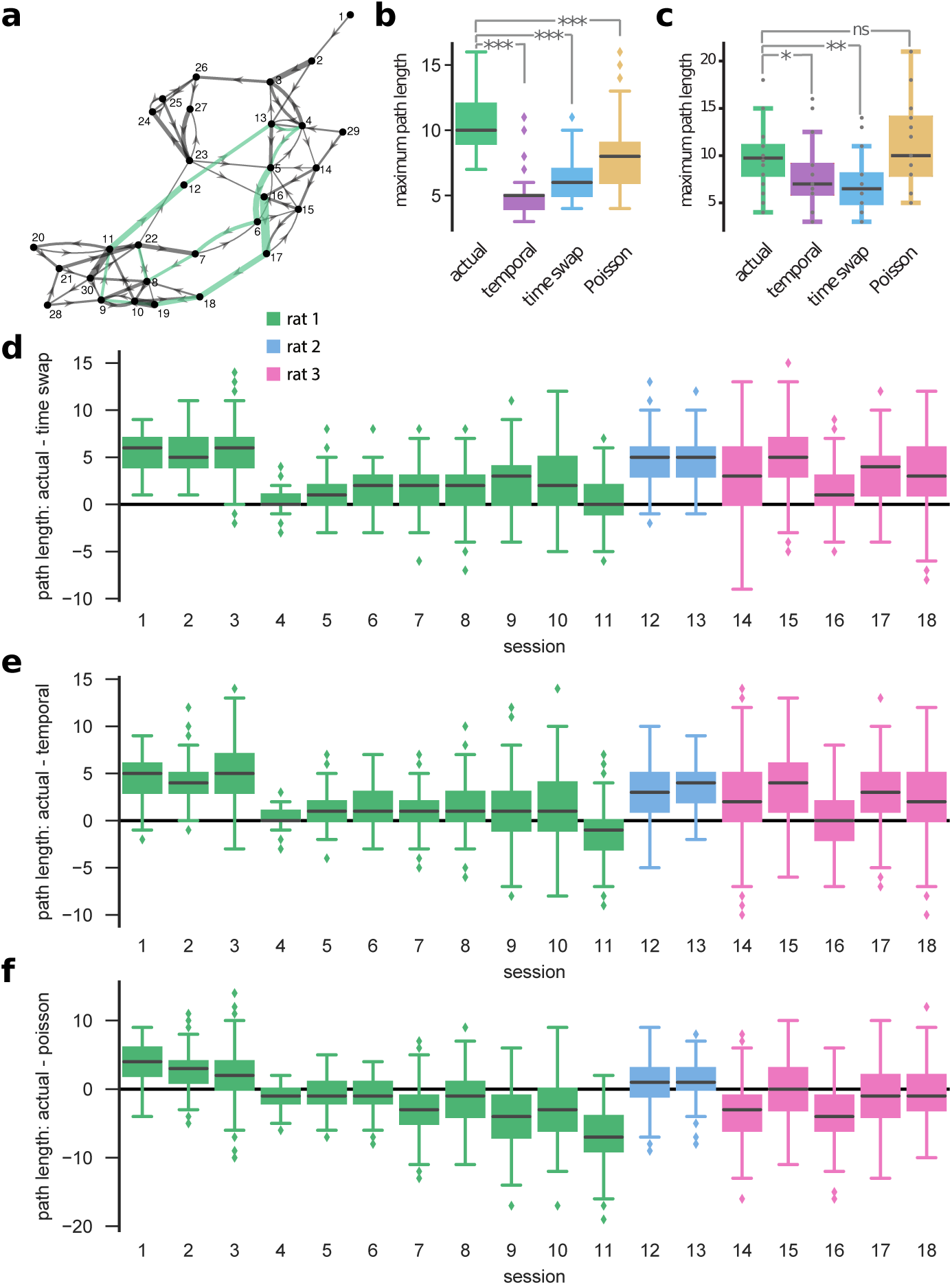
We calculated the longest path within an unweighted directed graph with the edges reflecting the transition probabilities (see Methods for details). **a.** The graph (force-directed layout) represents a model trained on actual data. For illustration purpose, we ignored the transition probabilities below 0.1 to reduce the clutter. The highlighted path shows the longest path in the example graph. **b.** For an example session, we computed the maximum path length for actual and corresponding shuffle datasets (temporal, time swap, and Poisson). **c.** The panel shows the Aggregate results built of median path lengths from all sessions. We see that the actual data on average result in longer sequences compared to time swap (*p* = 0.008, Mann-Whitney *U* test) and temporal surrogate datasets (*p* = 0.04, Mann-Whitney *U* test). On the contrary, no significant difference is seen when comparing with the Poisson datasets (*p* = 0.57, Mann-Whitney *U* test). Note that, however, due to non-sparseness of the observation matrix for a Poisson model (*Figure 2-Figure Supplement 2*), the long paths barely translate to sequences of transitions between distinct sparse co-activations. In panels **d-f**, difference between path lengths obtained from actual data and those from surrogate datasets are shown separately for all sessions : actual versus **d**. time swap, **e**. temporal, and **f**. Poisson. The actual data result in longer sequences compared to time swap and temporal shuffle datasets in most of the sessions (15 out of 18) (*p* < 0.001, Mann-Whitney *U* test). The Poisson data, however, result in sequences with similar (3 out of 18) or higher lengths (10 out of 18) compared to actual data.

**Figure 3-Figure supplement 1.**
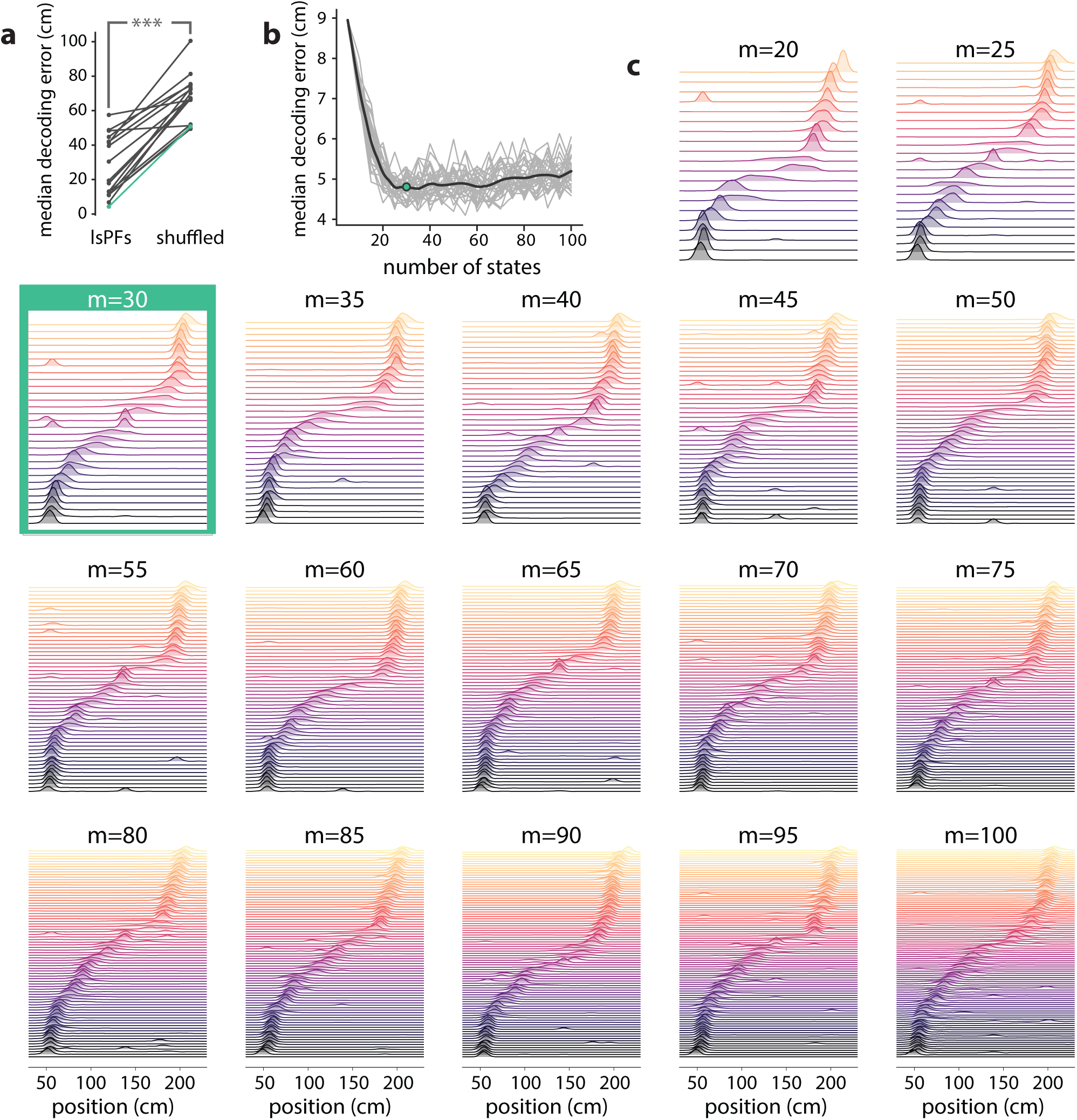
We investigated to what extent our PBE models encoded information related to the animal’s positional code by learning an additional mapping from the latent-state space to the animal’s position (resulting in a latent-space place field, lsPF), and then using this mapping, we decoded run epochs to position and assessed the decoding accuracy. **a.** We computed the median position decoding accuracy (via the latent space) for each session on the linear track (*n* = 18 sessions) using cross validation. In particular, we learned a PBE model for each session, and then using cross validation we learned the latent space to animal position mapping on a training set, and recorded the position decoding accuracy on the corresponding test set by first decoding to the state space using the PBE model, and then mapping the state space to the animal position using the lsPF learned on the training set. The position decoding accuracy was significantly greater than chance for each of the 18 sessions (*p* < 0.001, Wilcoxon signed-rank test). **b.** For an example session, we calculated the median decoding accuracy as we varied the number of states in our PBE model (*n* = 30 realizations per number of states considered). Gray curves show the individual realizations, and the black curve shows the mean decoding accuracy as a function of the number of states. The decoding accuracy is informative over a very wide range of number of states, and we chose *m* = 30 states for the analysis in the main text. **c.** For the same example session, we show the lsPFs for different numbers of states. The lsPFs are also informative over a wide range of number of states, suggesting that our analyses are largely insensitive to this particular parameter choice (the number of states).

**Figure 4-Figure supplement 1.**
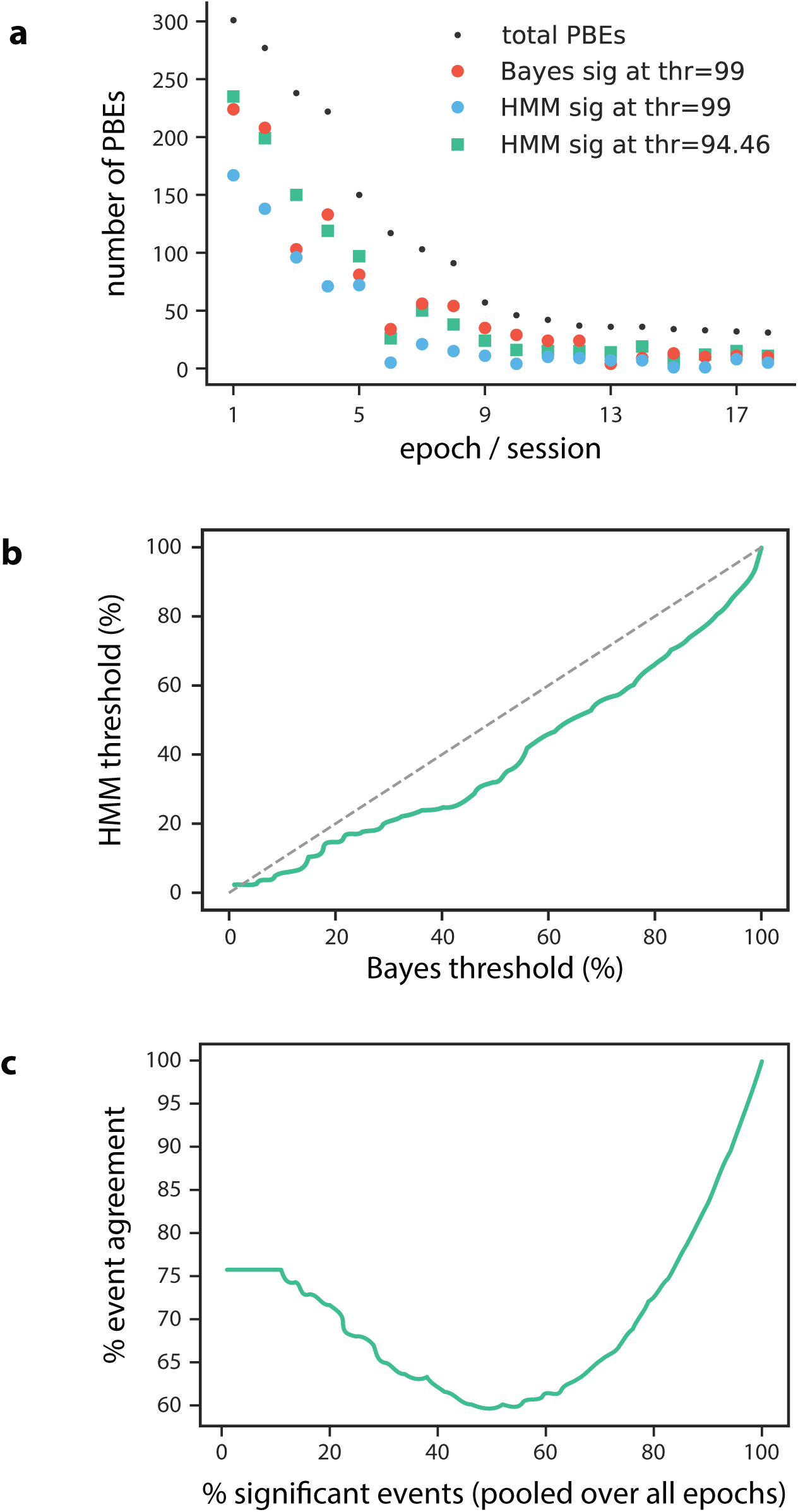
**a.** The number of Bayesian significant PBEs, as well as the total number of PBEs are shown for each session (*n* = 18) when using a significance threshold of 99%. We find that 57% of PBEs (1064 of 1883) are Bayesian significant at this threshold. When using this same threshold for the model-congruence (HMM) significance testing, we find that only 35% of PBEs (651 of 1883) are model congruent. In order to compare the Bayesian and model-congruence approaches more directly, we therefore lowered the model-congruence threshold to 94.46%, at which point both methods had the same number of significant events (1064 of 1883). **b.** For each Bayesian significance threshold, we can determine the corresponding model-congruence threshold that would result in the same number of significant PBEs. **c.** Using the thresholds from **b.** such that at each point, both Bayesian and model-congruence approaches have the same number of significant PBEs, we calculate the event agreement between the two approaches. We note that our chosen threshold of 57% significant events has among the worst agreement between the two approaches.

**Figure 5-Figure supplement 1.**
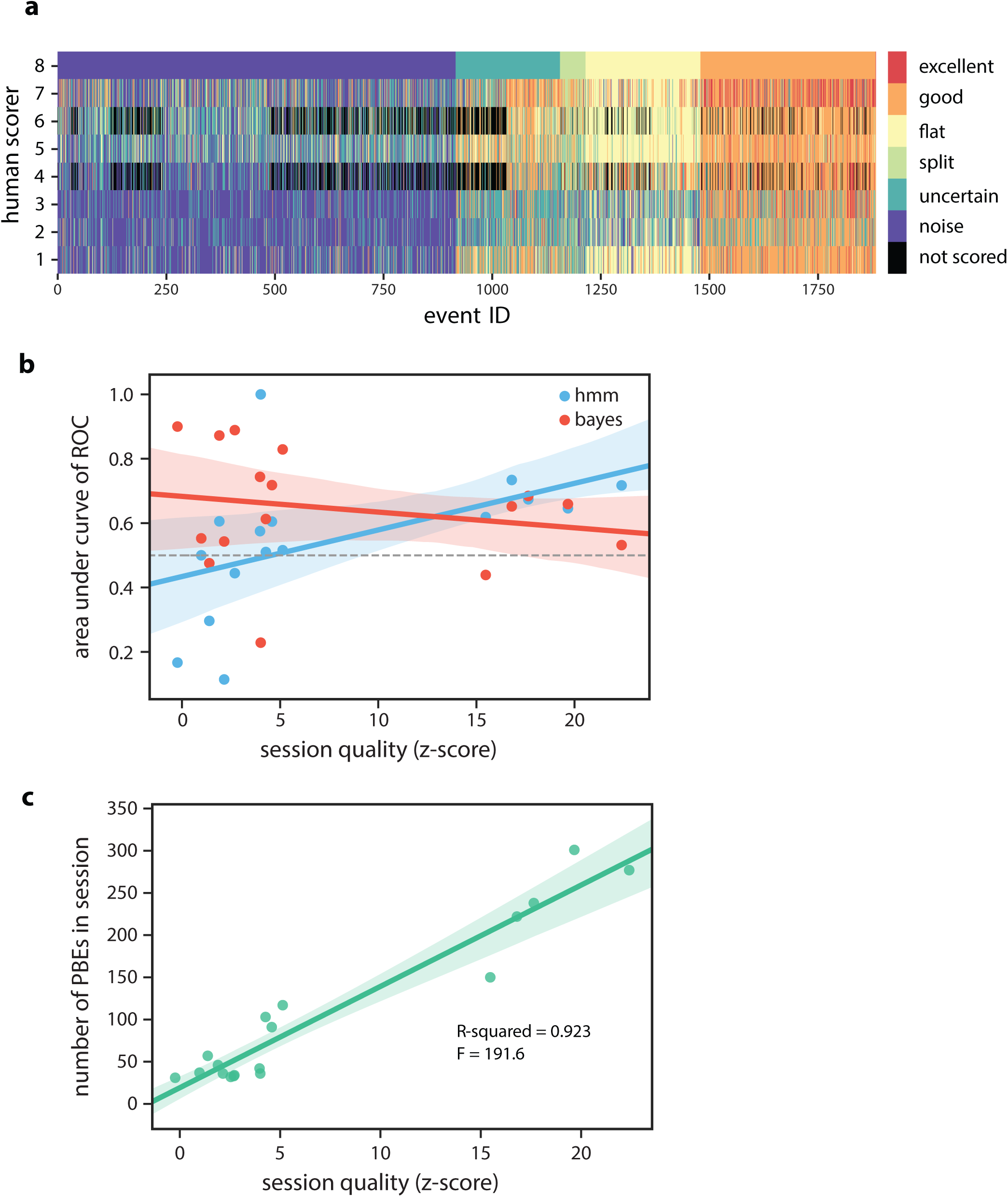
**a.** Manual scoring results from 8 human scorers (six individuals scored *n* = 1883 events, two individuals scored a subset of *n* = 1423 events). Events were presented to each participant in a randomized order, and individuals were allowed to go back to modify their results before submission. Here, events are ordered according to individual #8’s classifications. **b.** The model-congruence (HMM) approach appears to have higher accuracy when the session quality is higher (*R*^2^ = 0.26, *F* = 4.951), which is consistent with our expectation that we need many congruent events in the training set in order to learn a consistent and meaningful model. **c.** The session quality is strongly correlated with the number of PBEs recorded within a session (*R*^2^ = 0.92, *F* = 191.6).

**Figure 6-Figure supplement 1.**
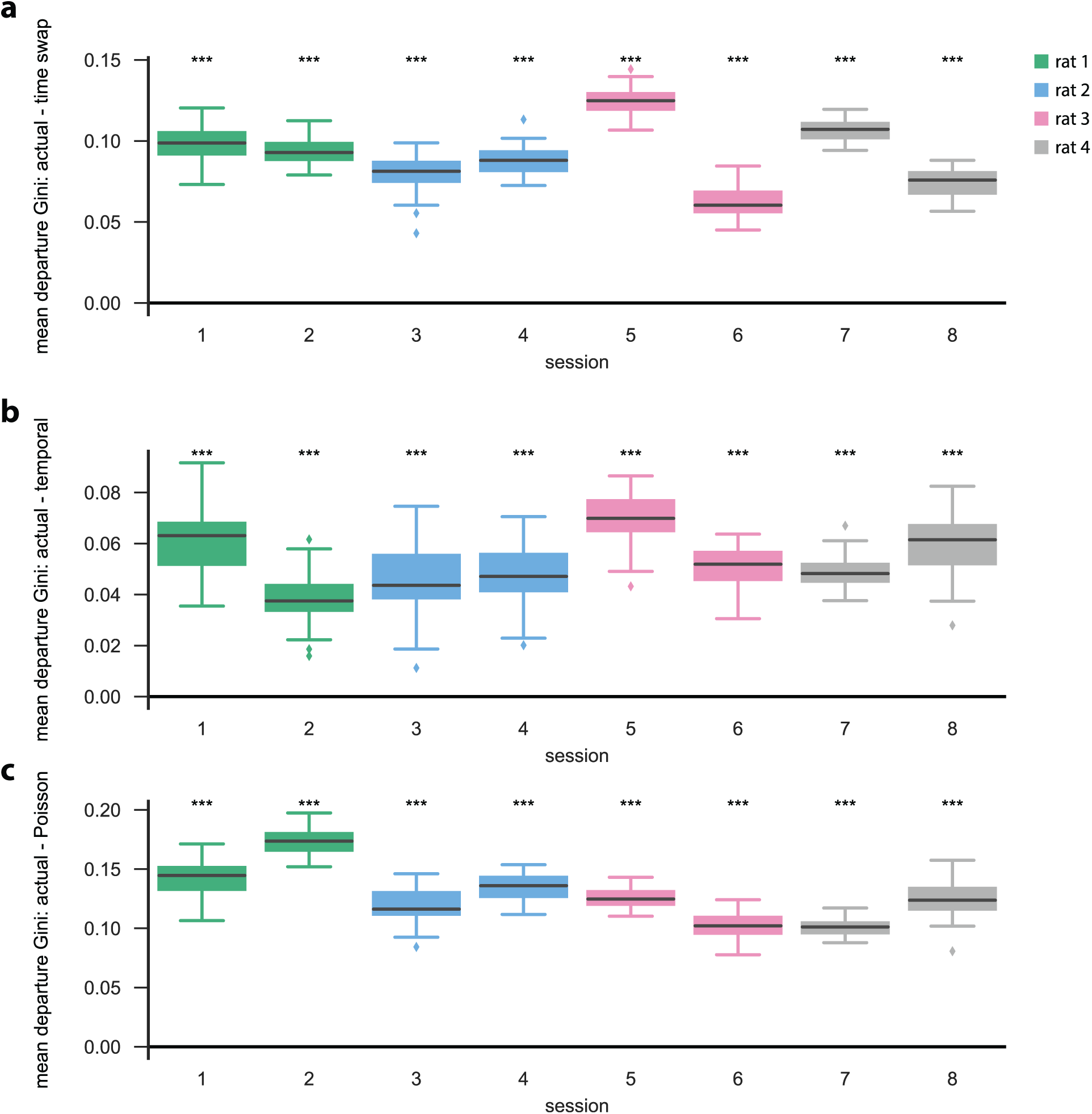
Similar to the linear track (one dimensional) case, we find that models learned on actual open field PBE data are significantly more sparse (here showing mean departure sparsity) than their shuffled (*m* = 50 shuffles) counterparts. This is true for each of the *n* = 8 open field sessions (*p* < 0.001, Mann-Whitney *U* test). **a.** Difference [in departure Gini coefficients] between actual and time swap test data, **b.** between actual and temporal test data, and **c.** between actual and Poisson surrogate data.

**Figure 6-Figure supplement 2.**
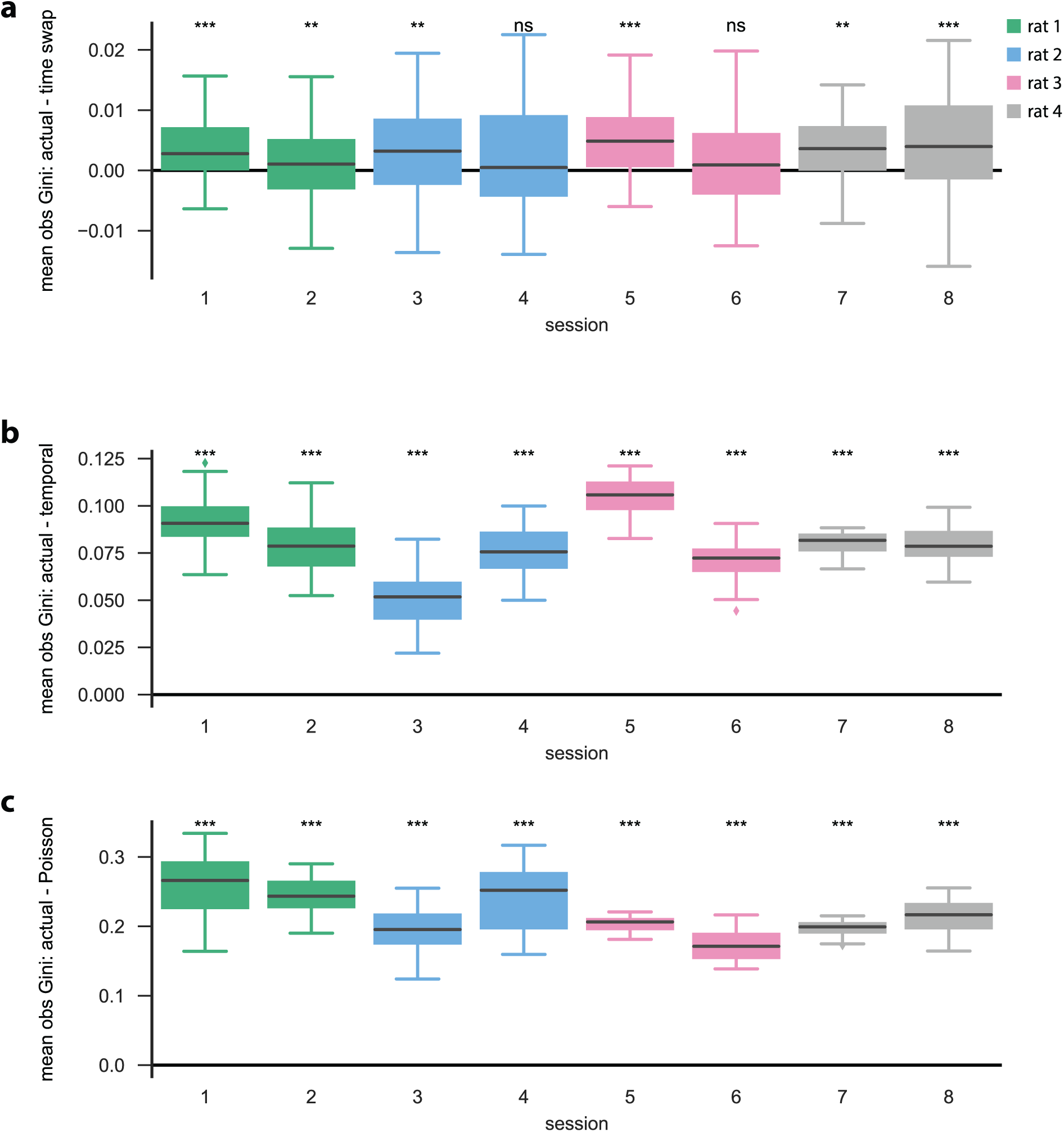
**a.** Difference [in observation sparsity Gini coefficients across states] between actual and time swap test data, **b.** between actual and temporal test data, and **c.** between actual and Poisson surrogate data. Similar to the linear track (one dimensional) case, we find that the observation sparsity across states for actual data are significantly greater than that of both the **b.** temporal and **c.** Poisson surrogates (for each session, *p* < 0.001, Mann-Whitney *U* test), and that **a.** for some sessions, there are no significant differences between the actual and time swap surrogates.

**Figure 6-Figure supplement 3.**
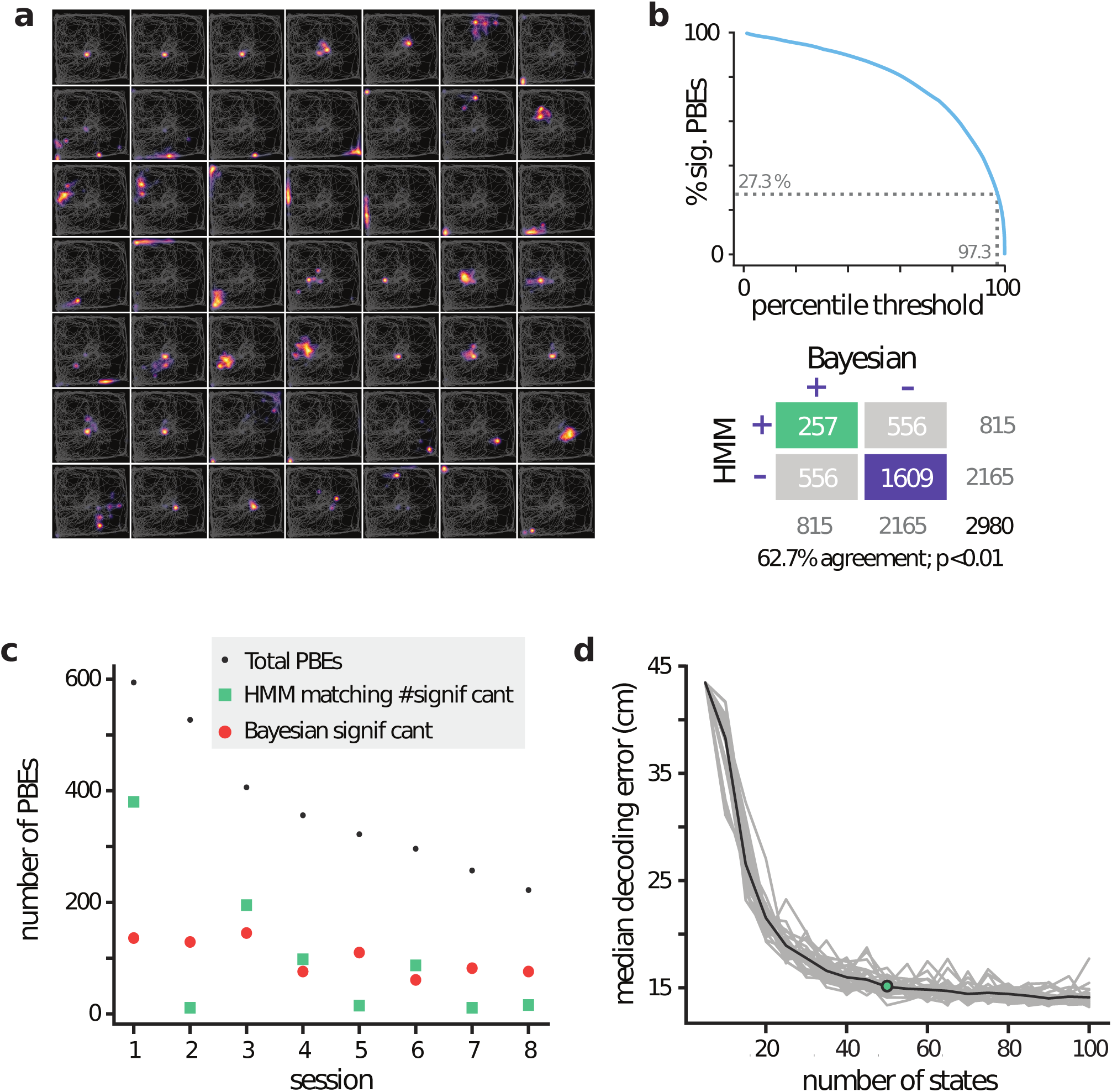
**a.** lsPFs for 49 of the 50 latent states from an example session. **b.** (Top) Effect of model-congruence threshold on the number of significant PBEs. (Bottom) Comparison matrix between Bayesian replay detection and our model-congruence approach, where the threshold was chosen to match the total number of significant events pooled overall 8 sessions. **c.** Comparison between number of significant Bayesian events vs number of significant events using our model-congruence approach, when choosing the threshold as in **b.**. Sessions are ordered in decreasing numbers of total PBEs. Note that session 1 is a significant outlier, causing mismatches between many other sessions (2, 5, 7, 8), suggesting that matching on a per-session basis may be more appropriate in this case. **d.** Median position decoding error (via the latent space and lsPFs) was evaluated using cross-validation in an example session (*n* = 30 realizations for each model considered, shown in gray, mean shown in black), indicating that (i) the PBE-learned latent space encodes underlying spatial information, and (ii) that our PBE models are informative about the underlying position over a wide range of numbers of states.

**Figure 6-Figure supplement 4.**
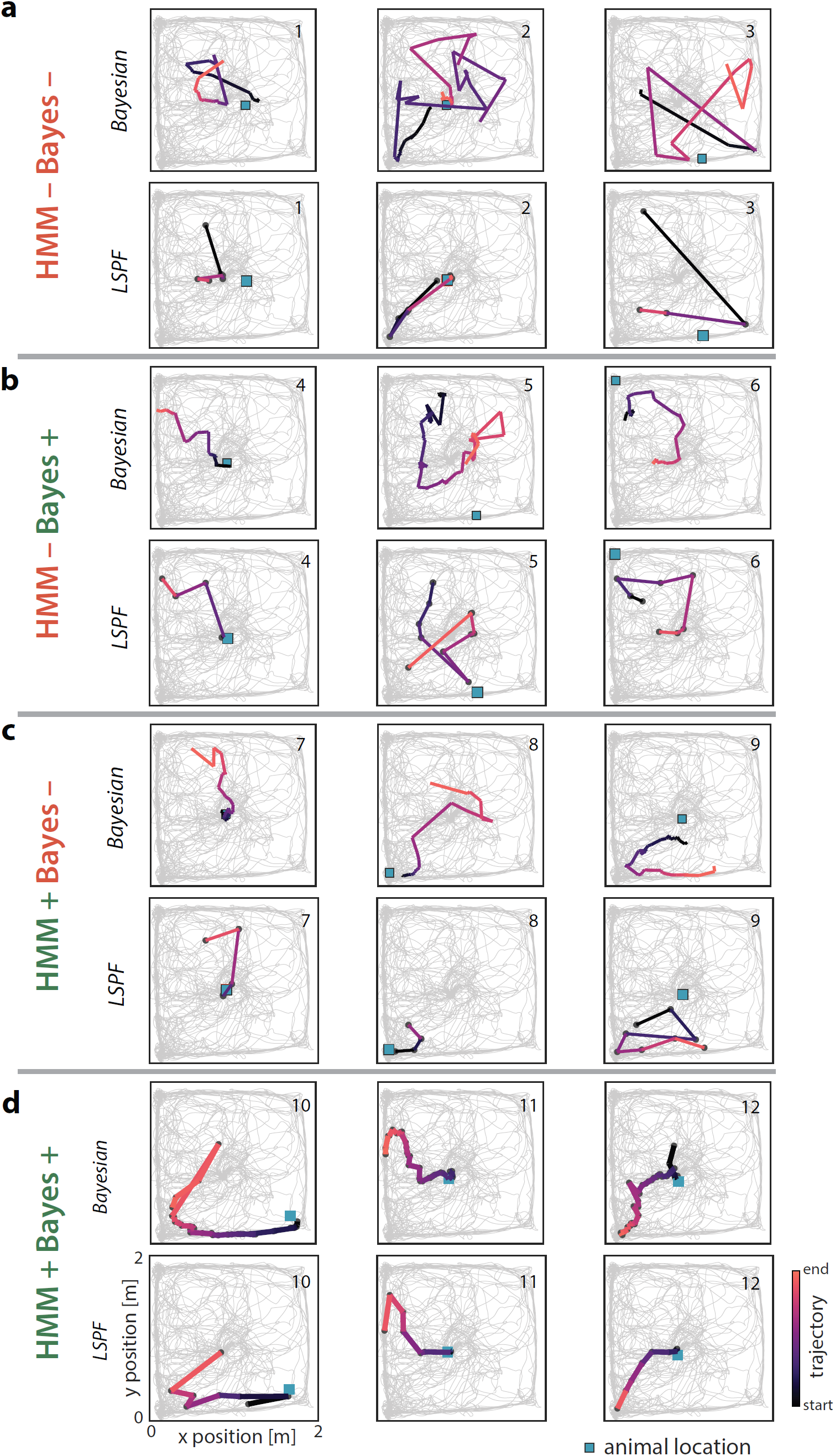
**a.** Three example PBEs are shown that were classified as non-significant by both the Bayesian and model-congruence approaches. The top row shows the PBEs decoded with place fields using a Bayesian decoder in 20 ms bins, with a 5 ms stride. The bottom row shows the same events, but decoded in 20 ms non-overlapping time bins using the lsPFs. **b.** Three example PBEs are shown that were classified as significant replay by the Bayesian approach, but not by the model-congruence approach. **c.** Three example PBEs are shown that were classified as significant replay by the model-congruence approach, but not by the Bayesian approach. **d.** Three example PBEs are shown that were classified as significant by both approaches.

